# A bacterial chaperone–usher pathway links interkingdom association to antimicrobial protection of a fungal partner

**DOI:** 10.64898/2026.10.29.762636

**Authors:** Alyssia Gonzalez, Kurt Dahlstrom

## Abstract

Antimicrobial compounds (AMCs) influence species-level composition of microbial communities, governing beneficial and pathogenic interactions with plant and mammalian hosts. Phenazine-1- carboxylic acid (PCA) is a bacterially produced, redox-active AMC abundant in the rhizosphere. A recent study discovered the PCA-susceptible fungus *Aspergillus calidoustus* can grow in PCA- abundant environments by associating with phenazine-tolerant bacteria, including the novel species *Paraburkholderia edwinii*. *P. edwinii’s* protection response is characterized by adherence to its fungal partner, bacterial aggregation, and sequestration and reduction of PCA, rendering PCA less toxic. Deletion of the bacterial transcriptional repressor HrcA generates a constitutively active protection response and enhanced protection of *A. calidoustus*. Here, we use the Δ*hrcA* strain to determine whether fungal protection uses bacterial self-protection machinery or dedicated interkingdom mechanisms. We identified a divergent archaic chaperone–usher pathway (CUP) operon encoding a predicted periplasmic chaperone, outer membrane usher, putative major pilin-like subunit, and putative adhesin-like tip protein. While many archaic CUP operons are annotated in diverse bacteria, their functions remain largely unexplored. We show these **f**ungal **a**ttachment **c**omplex (*fac*) genes promote bacterial-fungal attachment, are required for *P. edwinii* to sequester and detoxify PCA, and contribute to full fungal protection from PCA stress, despite not being required for *P. edwinii*’s own tolerance of PCA. We further identified the transcription regulator, FacR, that influences expression of the *fac* system. Understanding the functions of divergent archaic CUP systems may illuminate protective interkingdom partnerships and inform how pathogenic and beneficial microbes gain entry to AMC-replete environments, shaping microbial communities in agricultural and clinical contexts.

**IMPORTANCE:** Some species of fungi can be opportunistic pathogens, while others promote rhizosphere stability. Antimicrobial compounds help govern the composition of microbial communities by excluding susceptible organisms and selecting for microbes with intrinsic resistance. However, susceptible fungi can gain entry into microbial communities containing antagonist compounds by partnering with protective bacteria, and it is unknown whether bacteria are extending the mechanisms of self-tolerance to fungal partners, or if there is dedicated machinery for interkingdom protection. Here, we show that *Paraburkholderia edwinii* uses a divergent chaperone–usher system to associate with its fungal partner, *Aspergillus calidoustus*, and protect it from phenazine stress. This system promotes fungal attachment, antimicrobial sequestration, and maintenance of a reducing environment, yet is dispensable for the bacterium’s own survival. These findings demonstrate that bacteria can deploy dedicated machinery to protect partners across kingdoms, revealing that antimicrobial tolerance can emerge from microbial interactions rather than intrinsic resistance alone.

## INTRODUCTION

In nature, microorganisms seldom exist in isolation. Instead, they live in dynamic multi-species communities. These communities have a profound impact on the environments they reside in, including health outcomes in mammalian and plant hosts (1–5). The human immune system is modulated by the gut microbiota, impacting conditions such as irritable bowel disease (2), while the species-level composition of polymicrobial infections in cystic fibrosis patients play a large role in disease progression (5). Further, the presence of mycorrhizal and pathogenic fungi in the rhizosphere profoundly impacts crop health and yield (6, 7). These microbial communities’ functional outputs hinge on their species-level composition (8, 9). However, our knowledge of the impact specific microbial communities impart on their hosts and environments far outpaces our knowledge of how these communities are shaped for species-level membership in the first place. Thus, defining the molecular mechanisms involved in establishing and maintaining the species-level composition of polymicrobial communities is an important step in understanding and manipulating the positive and negative impacts they have on their hosts and environments.

One of the strongest leads into how microbes manage community membership is that microbes produce natural antimicrobial compounds (AMCs) which have been shown to mediate species-level composition of the communities in which they are produced (10). For example, phenazines are a class of small redox-active AMCs produced by many bacterial genera, including *Burkholderia*, *Streptomyces*, and most notably *Pseudomonas* (11, 12). Most bacterial phenazine biosynthetic pathways begin with the precursor molecule, phenazine-1-carboxylic acid (PCA), making PCA one of the most abundant AMCs in the natural environment (13, 14). PCA cycles through oxidized and reduced states, destabilizing metabolism and producing reactive oxygen species (ROS) that cause cellular damage (15). Historically, this has been understood to help shape microbial communities based on which microbes are sensitive to phenazine stress and which are tolerant. For example, many phenazine-tolerant *Paraburkholderia* spp. have important roles in plant growth promotion by enhancing nutrient acquisition and abiotic stress tolerance in communities where phenazine producers are present such as the rhizosphere (16). Conversely, several studies show that phenazine producers have an antagonist relationship with many important phenazine sensitive fungal species, including the human opportunistic pathogens *Aspergillus fumigatus* and *Candida albicans*, as well as phytopathogens such as *Fusarium oxysporum* (15, 17, 18). Surprisingly, phenazine-susceptible fungal species are frequently co-isolated with phenazine producing *Pseudomonas spp*. in environments such as the lungs of cystic fibrosis patients and the rhizosphere of dryland cereals (19, 20). Bacterial phenazine producers and phenazine sensitive fungi in mucosal lung model coinfections are even synergistically virulent, influencing severe disease progression (21). Bypassing phenazine stress by fungi that demonstrate inherent phenazine susceptibility is therefore an important topic for understanding how both pathogenic and beneficial fungi become fixed as members of a microbial community in AMC replete conditions.

A recent hypothesis addressing this question is that phenazine-tolerant bacteria can protect phenazine-sensitive organisms, such as desirable fungal species, in the rhizosphere. For example, a 2022 study discovered the phenazine-susceptible fungus, *Aspergillus calidoustus* can overcome PCA toxicity when paired with a native physically-associated partner, a novel phenazine-tolerant bacterium, *Paraburkholderia edwinii* (22). This revealed a microbe-microbe protective partnership that plausibly underpins the frequent co-isolation of phenazine susceptible fungi and phenazine producers, offering one mechanism by which the species-level composition of microbial communities can be shaped via otherwise broadly-acting AMCs.

Activation of the bacterial protection program of *A. calidoustus* from PCA stress includes four primary characteristics including adherence to the fungus, the formation of mature bacterial aggregates within the fungal mycelium in response to PCA exposure, acting as a ‘toxin sponge’ as *P. edwinii* sequesters PCA, and creating a microenvironment with a low reduction potential (22). Therefore, fungal PCA stress is managed via bacterial sequestration of PCA away from its fungal partner, and detoxifying PCA by limiting the molecule from cycling its redox state, thus preventing ROS production. Critically, this protective microbe-microbe interaction is not unique to *P. edwinii* and *A. calidoustus*. Several members of the *Paraburkholderia* genus have been demonstrated to protect fungi when paired together in the lab environment, including offering protection to the human pathogen *A. fumigatus* and phytopathogen *Penicillium glabrum* (22). Additionally, physically associated bacteria isolated from the environment have been shown to offer protection to fungi from diverse AMCs (23). These previous studies highlight how microbial partnerships may be a common mechanism to allow otherwise susceptible organisms to be selected to enter AMC-replete environments and emphasize the importance in developing an understanding of the molecular mechanisms bacteria employ during these protective microbe-microbe interactions.

In our present study, we identify bacterial genetic elements required for the protection program by conducting an RNA-seq analysis on a strain of *P. edwinii* in which the protection response is locked in the active state. HrcA is a transcriptional repressor previously identified as having a role in regulating the protection program (22). When deleted, the Δ*hrcA* strain promotes more fungal growth in the presence of PCA than wildtype, sequesters significantly more PCA, has an enlarged reducing zone, and forms enhanced aggregates in fungal mycelium even in the absence of PCA, indicating the protection program is always active (22). By comparing RNA transcripts in the mutant to the wild-type strain, we identified a divergent archaic chaperone–usher pathway (CUP) operon that encodes putative pilus-like adhesins and contributes to the protective partnership between *P. edwinii* and *A. calidoustus*. These **f**ungal **a**ttachment **c**omplex (*fac*) genes encode the putative major pilin subunit, FacA, a putative periplasmic chaperone, FacB, a putative outer membrane usher, FacD, and a putative adhesive tip, FacG.

CUP operons are widespread in bacteria. They are classified by the amino acid sequence of their outer membrane usher protein into three broad groups that contain six phylogenetic clades: Alternative (α-fimbriae), Classical (β-fimbriae, γ-fimbriae, κ-fimbriae, π-fimbriae) and Archaic (σ-fimbriae) (24). CUP operons are primarily known to encode pili or fimbriae structures. However, a couple atypical CUP operons have been identified to encode alternative extracellular products. For example, *Yersinia pestis’s caf1* γ-clade CUP secretes a gel-like F1 capsule that helps the bacterium resist phagocytosis (25), while *Myxococcus xanthus’s* MXAN_3885-2 archaic CUP operon secretes a spore-coat protein during spore formation (24, 26). Most knowledge of CUP operon product biosynthesis and function comes from classical CUP fimbriae studies, such as Fim and Pap pili of *E. coli* (27, 28). Pili are extracellular filamentous structures anchored to the cell surface and are involved in adherence to biotic and abiotic surfaces and influence biofilm-dependent antimicrobial resistance (27, 29–33). Additionally, known atypical CUP operons share this self-protection function by producing capsule or spore coat to protect them from environmental stress (25, 26), therefore, making CUP operons a promising target to study a protective bacterial-fungal interaction, regardless of their extracellular product.

Archaic CUP operons are thought to be the most ancestral and widespread, with many non-canonical archaic systems being bioinformatically identified across diverse bacteria phyla (24). Further, archaic CUP are the least understood CUP systems with only the Csu pili of *Acinetobacter baumannii,* CupE pili of *P. aeruginosa* and MXAN_3885-2 system of *M. xanthus* being well studied representatives of this clade (34–37). Archaic CUP systems of these model organisms are implicated in self-protection, with MXAN_3885-2 system secreting protective spore coat and the super-elastic archaic CUP pili participating in biofilm dependent antimicrobial resistance through self-adherence, abiotic adhesion, and epithelial cell adhesion. However, these functions are described from only a few canonical archaic CUP systems and many non-canonical archaic CUP operon functions remain unknown (29, 35–38). Given the diversity in physical features and functions of CUP operons, we therefore explored whether the non-canonical archaic CUP operon that is upregulated during *P. edwinii*’s protection response is responsible for self-protection, fungal partner-protection, or both.

Here, we demonstrate that loss of the *fac* genes produces a significant defect in fungal protection from PCA stress, while having no effect on *P. edwinii’s* own survival on PCA or self-adherence in the formation of bacterial aggregates, revealing a previously unrecognized role for archaic CUP systems. Specifically, *fac* deletion mutants showed a defect in adhering to fungal mycelium and were less efficient at sequestering and detoxifying PCA than wild-type, accounting for three of the four major bacterial protection behaviors governed by HrcA. Therefore, *P. edwinii* uses its non-canonical archaic CUz system to associate with its fungal partner and execute key components of the protection program in response to PCA. Finally, we identified an AraC-family transcriptional regulator, FacR, that promotes the expression of the *fac* operon in *P. edwinii* in response to PCA. Taken together, these results demonstrate that archaic CUP systems can be activated by AMCs relevant to microbial community membership, play an important role in associating with fungal hosts, and help extend AMC protection to other domains of life. This work expands the function of non-canonical archaic CUP systems beyond self-protection and begins to define the mechanisms by which microbes can enter into protective relationships to promote AMC resistance within beneficial microbial communities such as the rhizosphere as well as within sites of polymicrobial infections (22, 24, 39).

## RESULTS

### Components of an archaic chaperone-usher pathway operon are upregulated in a *P. edwinii* genetic background that locks the protection response in the active state

A previous study identified the bacterial transcriptional repressor HrcA as involved in *P. edwinii’s*protection response (22). This involvement was determined by the enhanced protection the Δ*hrcA P. edwinii* strain provides *A. calidoustus* against PCA stress compared to the wild-type strain, and the protection program remaining active even in the absence of PCA (22) (Fig. 1A). The enhanced protection phenotype includes four features: enhanced adherence in co-colony, enhanced bacterial aggregation within the mycelium, enhanced sequestration of PCA, and an enlarged reducing zone for PCA detoxification (22).

**Figure 1:**
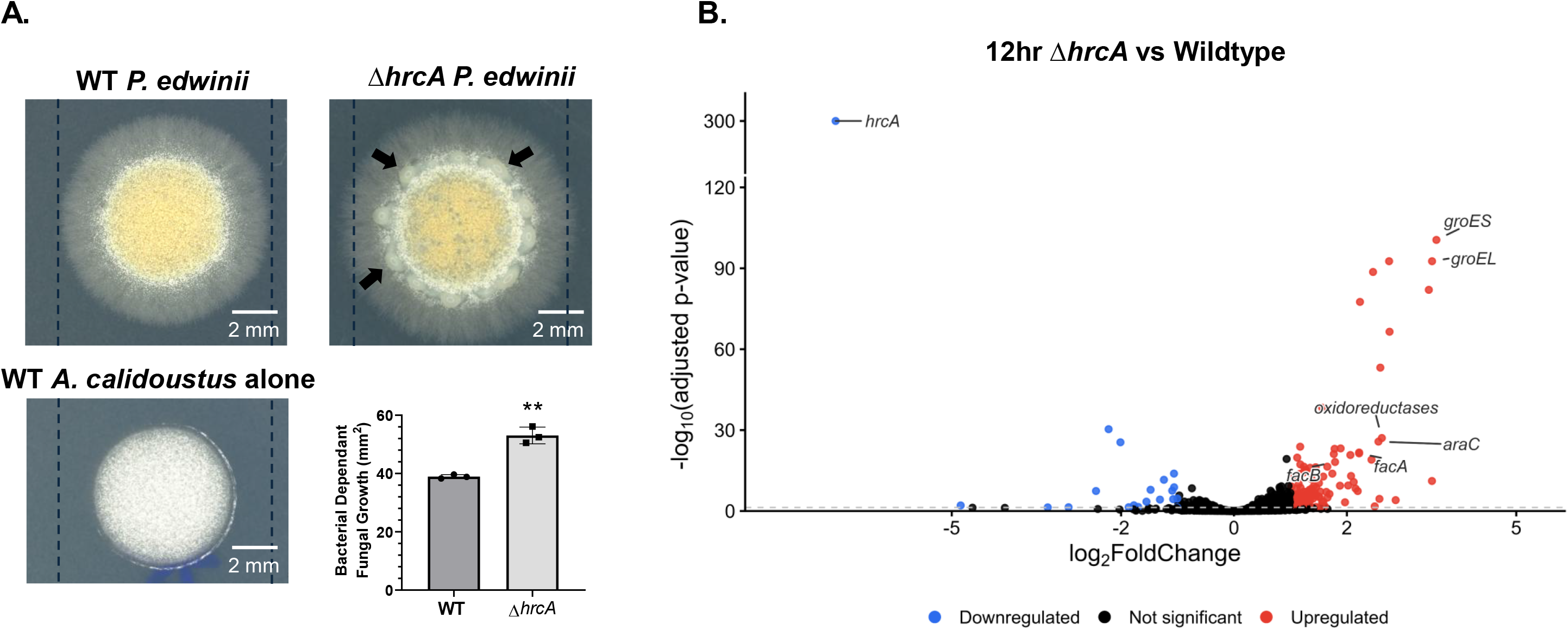
The first two genes of a Chaperone-Usher Pathway (CUP) and an AraC-family transcriptional regulator are upregulated in a *P. edwinii* strain that’s protection response is locked in the active state. (A) Wild-type and Δ*hrcA P. edwinii* in co-colony with *A. calidoustus,* and *A. calidoustus* alone on PDA containing 300µM PCA. The *P. edwinii* transcriptional repressor deletion strain, Δ*hrcA*, increases bacterial dependent fungal growth on 300µM PCA and has enhanced bacterial aggregation (black arrows) compared to wild-type. Represents three biological replicates. Scale bar indicates 2 mm, and error bars indicate standard deviation. Dashed lines indicate diameter of wild-type co-colony. Y-axis indicates bacterial-dependent fungal growth, with zero indicating the average area of three biological replicates of *A. calidoustus* grown alone on PCA. **p value<0.01. (B) Volcano plot showing the differential gene expression between Δ*hrcA and* wild-type *P. edwinii* on PDA for 12 hrs. Represents three biological replicates. Red transcripts are up-regulated and blue transcripts are down-regulated in Δ*hrcA* strain. Horizontal dash line indicated 0.05 adjusted p-value.

To identify candidate bacterial genes involved in these elements of fungal protection against PCA, we conducted an axenic RNA-seq to compare the differential gene expression between the Δ*hrcA* and wild-type strains of *P. edwinii* in the absence of PCA, where only the Δ*hrcA* strain’s protection response is active. Given that homologues of HrcA have been characterized as transcriptional repressors (40), we looked for any genes that were derepressed or otherwise upregulated in this background. The RNA-seq revealed 109 genes were significantly upregulated at 12 hours in the Δ*hrcA* background, falling into three broad functional groups (Supplementary Table 1). The first and most significantly upregulated group contained genes that encode the GroE protein folding chaperone complex, *groEL* (KZJ38_16355) and *groES* (KZJ38_16360) (Fig. 1B and Supplementary Fig. 1). The GroE complex is essential to most bacterial species (41), but *P. edwinii*’s genome contains two copies of these genes, and the deletion of the *groEL* locus upregulated in the RNA-seq did not yield an obvious fungal protection defect (Supplementary Fig. 2). However, *groEL* and *groES* are bona fide targets of the stress-inducible transcriptional regulator HrcA homologue found in *Bacillus subtilis*, supporting our RNA-seq results (40, 42). The RNA-seq also revealed a second group of upregulated genes with annotations that suggest functions in central metabolism that may be involved in regulating the cell’s internal redox state under PCA stress, including an NADP-dependent oxidoreductase, α/β hydrolases, and a cytochrome P460 family protein. However, most mutants in these metabolism-associated genes did not show an obvious defect in the *P. edwinii* protection response, and a protection defect observed for the NADP-dependent oxidoreductase mutant was not restored by plasmid-based complementation (Supplementary Fig. 2 and 3). The third and final group contained components of an archaic chaperone-usher pathway (CUP) operon we have named **f**ungal **a**ttachment **c**omplex (*fac*), including a putative major pilin-like subunit (*facA*) and a putative periplasmic chaperone (*facB*) that were upregulated in the Δ*hrcA* strain (Fig. 1B). Canonical archaic CUP pili have been implicated in self-protection, biofilm formation and epithelial host cell adhesion (35, 37). Many non-canonical archaic CUP operons have been bioinformatically identified in diverse bacterial phyla, yet their functions have not been defined. Therefore, we explored whether this non-canonical archaic CUP operon is involved in fungal protection through physical bacterial-fungal association, bacterial aggregation, PCA sequestration, and PCA detoxification during the protection response. Intriguingly, an AraC-family transcription factor was also found to be upregulated in the Δ*hrcA* strain (Fig. 1B). CUP encoding operons have previously been shown to be regulated by AraC-family transcription factors, suggesting similar regulatory mechanisms may be operating in *P. edwinii* (43), which we explore below.

The first two genes of *P. edwinii’s* CUP operon, *facA* and *facB,* are followed by a gene encoding an outer membrane usher protein (*facD*) and a putative adhesive tip subunit (*facG*) (Fig. 2A). Both FacA and FacG proteins contain the conserved domain, COG5430, that is also found in Csu pilin subunits and spore coat proteins (26, 34). The position of the *facG* at the distal region of the operon suggests it may be functioning as an adhesive tip protein, such as the case with *CsuE* of *A. baumannii* CUP pili (30). Additionally, the predicted structure of FacG contains a hydrophobic pocket with two tyrosine and a hydrogen bond forming residue (Asp), known features of the adhesion *fimH* of *E. coli* CUP pili (Fig. 2B and Supplementary Fig. 4) (44). Moreover, FacG has a positive electrostatic patch that may be involved in electrostatic steering towards the negatively charged fungal cell wall (Fig. 2B). Considering the *fac* system’s subunits shares the COG5430 domain with Csu pilin subunits we assessed whether FacA and FacG are structurally compatible with a pilus-like assembly. We used SignalP to identify predicted N-terminal signal peptides, which were removed prior to structural modeling and AlphaFold3 (45) to model a putative complex containing two copies of the predicted mature FacA protein and one copy of the predicted mature FacG protein (Fig. 2A and Supplementary Fig. 4). AlphaFold3 predicted an elongated arrangement with apparent FacA–FacA and FacA–FacG contacts. The model had a pTM score of 0.56 and an ipTM score of 0.44 (Fig. 2A and Supplementary Fig. 4). While this indicates limited confidence in the overall static architecture and interchain interfaces, these scores could also suggest an elastic and dynamic nature of an archaic CUP pilus. Thus, the results are consistent with structural compatibility between FacA and FacG in a pilus-like assembly model, but do not rule out the possibility of an alternative extracellular product.

**Figure 2:**
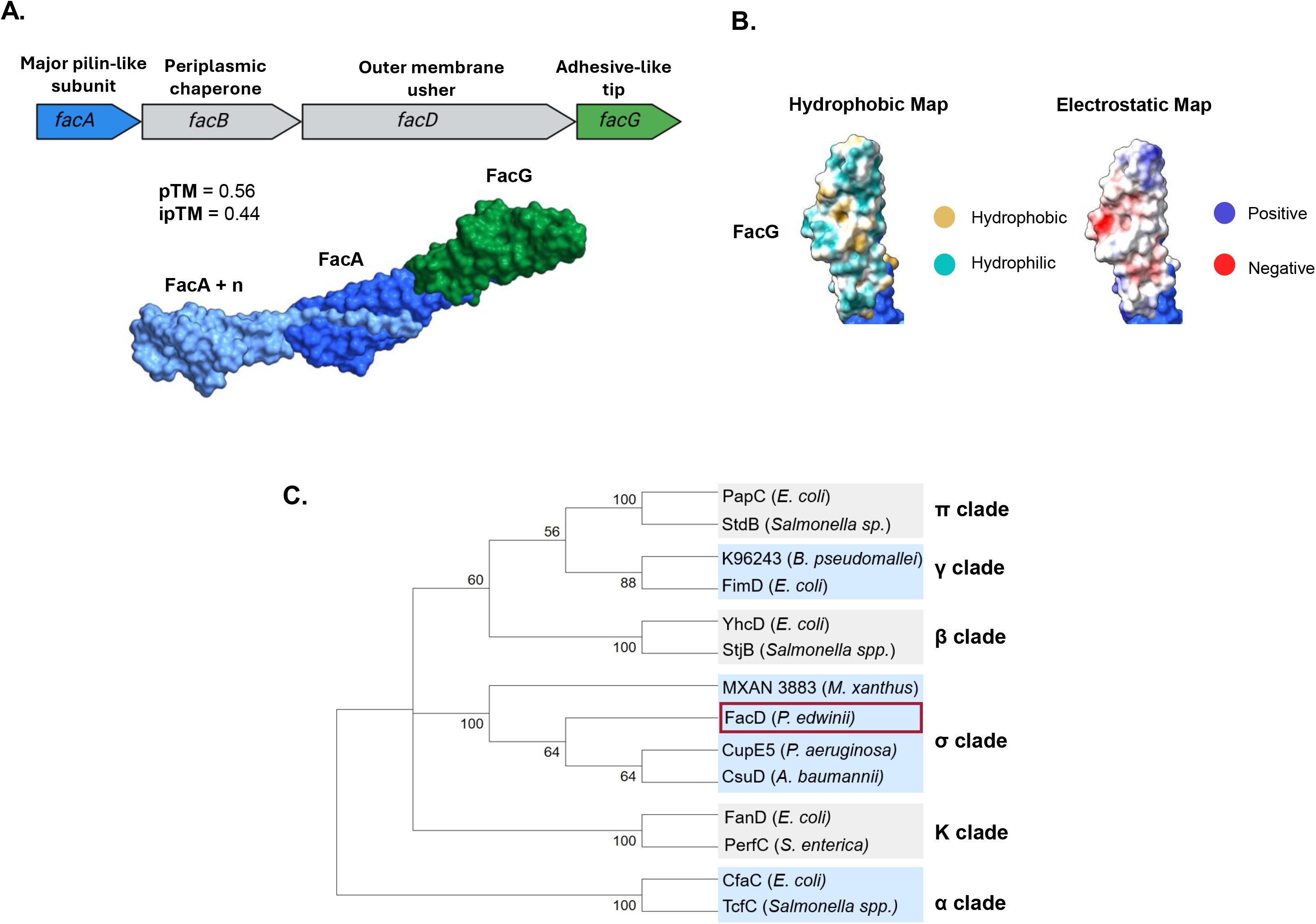
The *fac* system is structurally compatible with a pilus-like assembly and falls under the archaic clade of CUP. (A) Schematic of Chaperone-Usher Pathway (CUP) operon, *fac*. Created in BioRender. Dahlstrom, K. (2026) https://BioRender.com/qcxhkis. Alphafold3 predicted polymer containing two copies of mature FacA and one copy of mature FacG. pTM is the overall fold/topology prediction score (0-1) and the ipTM is the interface prediction score (0-1). (B) Hydrophobic and electrostatic maps of FacG, yellow is hydrophobic, cyan is hydrophilic, red is negative and dark blue is positive electrostatic. (C) Cladogram of CUP outer membrane usher protein amino acid sequences to classify *P. edwinii’s fac* operon. Values at nodes are bootstrap consensus values (0-100).

Archaic clade (σ) of CUP are the most prevalent yet understudied CUP clades (24). Therefore, to validate that the *fac* operon falls into the archaic CUP clade, we constructed a cladogram using the amino acid sequence of the outer membrane usher proteins, placing *P. edwinii’s* outer membrane usher protein, FacD, within the archaic clade (σ) with a consensus bootstrap value of 100 (Fig. 2C) (24). FacD was grouped as more closely related to CsuD and CupE5, archaic usher proteins that secrete Csu pili of *A. baumannii* and CupE pili of *P. aeurginosa* respectively, than to MXAN 3883 of *M. xanthus* which secretes spore-coat protein, with a consensus bootstrap value of 64 (26).

### The *fac* system contributes to *P. edwinii’s* protection of *A. calidoustus* from PCA stress but is not necessary for *P. edwinii’s* own survival

Given the non-canonical nature of this archaic CUP operon and its upregulation in the Δ*hrcA* background, we tested whether these genes contribute to phenotypes expected for a CUP-associated adhesin system and to any of the four protection characteristics of the Δ*hrcA* strain discussed above. First, to determine if the *fac* system is involved in *P. edwinii’s* protection of *A. calidoustus* from PCA assault, we made in-frame deletions of each individual component as well as the entire *fac* operon. These mutant strains were evaluated for primary protection defects in the presence of the fungus by quantifying their bacterial-dependent fungal growth area in the presence of PCA. Each *fac* component single deletion, as well as the whole operon deletion, had a significant protection defect compared to the wild-type strain of *P. edwinii* (Fig. 3A). Although CUP components are historically difficult to complement due to their tightly ordered assembly that relies heavily on stoichiometry balancing (31, 43, 46, 47), we were able to complement the *fac* operon and partially complement *facG* from a plasmid using an inducible promoter (Fig. 3B). These results indicate that *P. edwinii’s fac* system plays a role in the protection of *A. calidoustus* from PCA stress.

**Figure 3:**
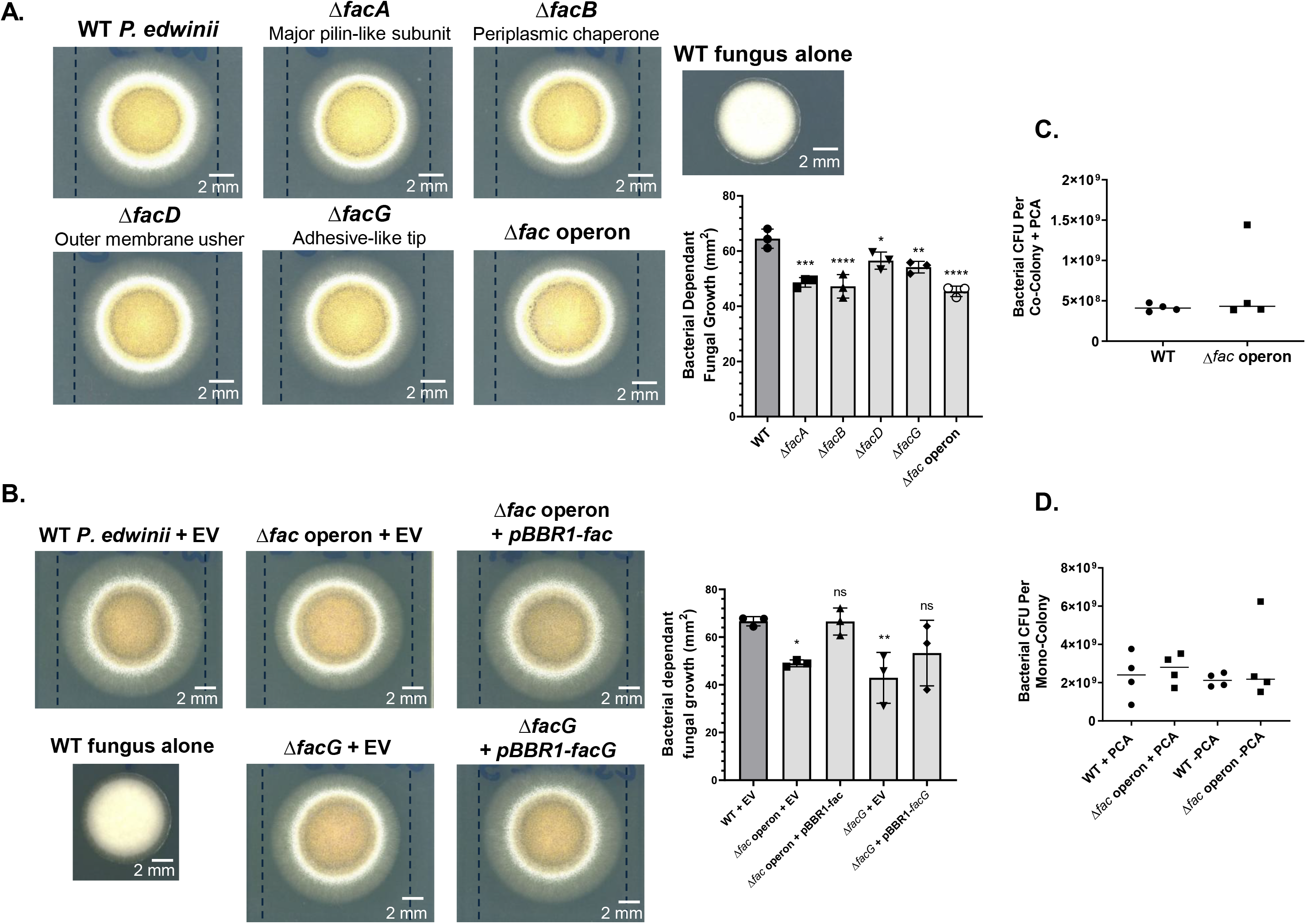
The *fac* system contributes to *P. edwinii’s* protection of *A. calidoustus* from PCA stress but is not necessary for *P. edwinii’s* own survival. (A-B) Since plasmid carriage impacts *P. edwinii* growth under these conditions, deletion mutants were compared to plasmid-free wild-type at 68 hrs, whereas complemented strains were compared to the corresponding wild-type empty vector (EV) control at 90 hrs. Represents three biological replicates. Scale bar indicates 2 mm, and error bars indicate standard deviation. Y-axis indicates bacterial-dependent fungal growth, with zero indicating the average area of three biological replicates of *A. calidoustus* grown alone on PCA. *p value<0.05, **p value<0.01, ***p value<0.001, ****p value<0.0001. (A) *fac* system mutants in co-colony display a statically significant decrease in bacterial dependent fungal growth on 300µM PCA compared to wild-type. Dashed lines indicate diameter of wild-type co-colony. (B) *fac* operon and *facG* complement strains in co-colony grown on PDA with 300µM PCA and 250µM L-rhamnose. Gene expression from pBBR1 under a rhamnose inducible promoter. Dashed lines indicate diameter of wild-type + EV co-colony. (C) Δ*fac* operon bacterial CFU in co-colony on 300µM PCA compared to wild-type. Represents four biological replicates. (D) Δ*fac* operon bacterial CFU in mono-colony in the presence and absence of 300µM PCA compared to wild-type. Represents four biological replicates.

In protective bacterial-fungal partnerships, it is unknown whether the bacterium is extending processes it uses for self-protection from AMCs to the fungal partner, or if the bacterium has machinery which may specifically enable fungal survival from AMCs. Importantly, the results from Figure 1B could be explained either by *P. edwinii* requiring the *fac* system to enable fungal survival when treated with PCA, or by the *fac* system being required for *P. edwinii*’s own survival against PCA. To determine if the *fac* mutants had a survival defect in protection response activating conditions, we quantified the bacterial colony forming units (CFUs) from the bacterial-fungal co-colony when treated with PCA (Fig. 3C). The Δ*fac* operon strain had comparable numbers of bacterial CFUs as the wild-type, indicating the *fac* system is not necessary for *P. edwinii’s* survival in co-colony when treated with PCA. To validate that the deletion of the *fac* operon does not impact bacterial survival in general, we quantified bacterial CFUs in monoculture grown in the presence and absence of PCA. Again, there were comparable numbers of bacterial CFUs across wild-type and Δ*fac* operon strains grown under either condition (Fig. 3D). These results indicate the *fac* system is not involved in self-protection against PCA stress, and instead it is specific to interkingdom protection of the fungal partner and thus contributes to the enhanced protection provided by the Δ*hrcA* strain.

### The *fac* system promotes bacterial adherence to *A. calidoustus* mycelium and formation of micro-aggregates upon initial bacterial-fungal association

It has been documented that archaic CUP that secrete pili function in adhesion to biotic and abiotic surfaces (27, 46). In *Acinetobacter baumannii,* Csu pili have been described as poly-adhesive and have been shown to adhere to human host epithelial cells in a biofilm-like manner (29, 36). We hypothesized that *P. edwinii* may be using the *fac* system similarly, playing a role in fungal association in this model system. To test the *fac* system’s role in bacterial-fungal association, we developed an adhesion assay to quantify the number of bacterial CFUs adhered to fungal mycelium within the first 30 minutes of co-incubation in potato dextrose broth (PDB) (Fig. 4A). The Δ*fac* operon had a significant decrease in bacterial CFUs adhered to the *A. calidoustus* mycelium compared to wild-type, indicating the *fac* system plays a role in initial bacterial-fungal association in this model system (Fig. 4B). Interestingly, the deletion of Csu pili adhesive tip, *csuE*, in the *A. baumannii* 19606 strain has shown to allow for increased adhesion to epithelial cells that is thought to be caused by an unknown feedback loop (29, 48). The Δ*facG* mutant showed a less severe fungal adhesion defect compared to the Δ*fac* operon, reflecting adhesion results of *A. baumannii* 19606 archaic CUP pili. To understand if the *fac* system is the primary initial adherence system or if alternative adhesion systems will compensate for it overtime, we conducted the adhesion assay with a three-hour co-incubation time point, six times longer than the initial 30 mins. At three hours there is some adherence compensation occurring, however the Δ*fac* operon still displays a minor adherence defect (Supplemental Figure 5).

**Figure 4:**
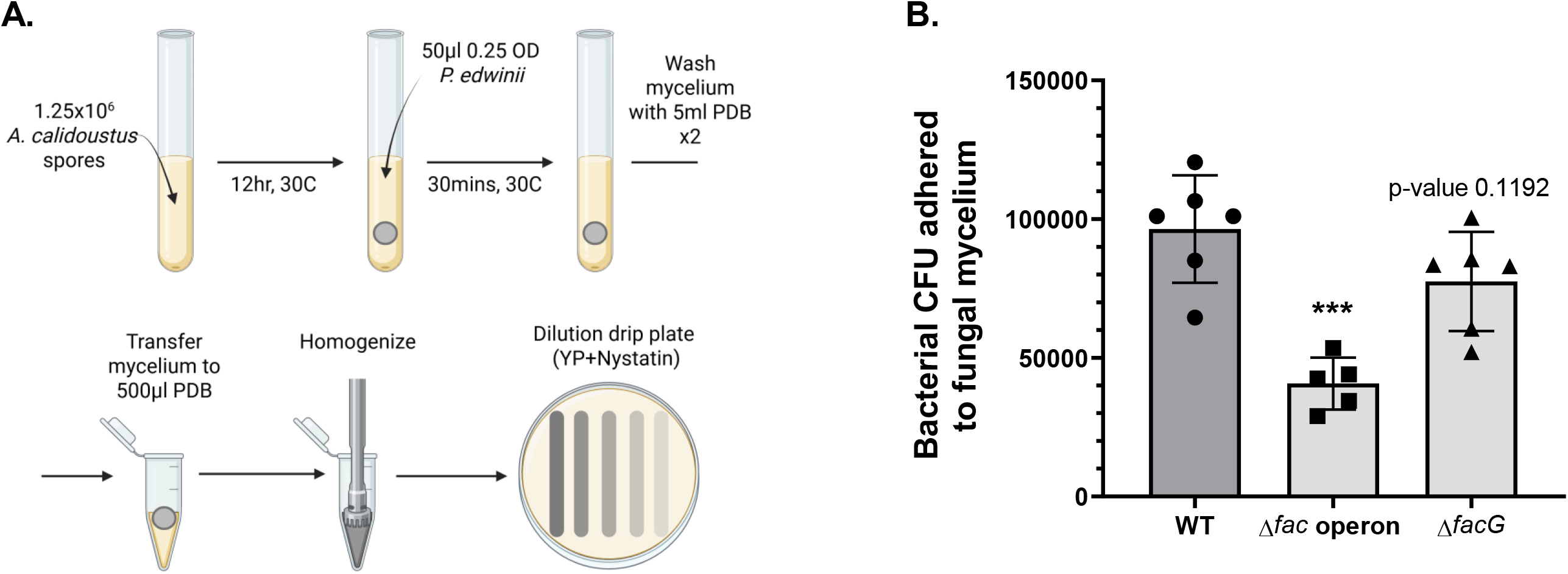
The *fac* system promotes bacterial adhesion to fungal mycelium during first 30 minutes of co-incubation. (A) Schematic of adhesion assay workflow. Created in BioRender. Dahlstrom, K. (2026) https://BioRender.com/xhrdn0v. (B) Wild-type, Δ*fac* operon, and Δ*facG* bacterial CFUs adhered to *A. calidoustus* mycelium after 30 minutes of co-incubation in 5ml PDB. Error bar is standard deviation. Represents six to five biological replicates. One replicate of Δ*fac* operon was lost during processing. ***p value<0.001.

Although the adhesion assay reveals the involvement of the *fac* system in bacterial-fungal association, the assay does not reveal what morphological changes are present during adherence to the fungal partner. To validate the adhesion assay results and gain resolution of *fac* system function during physical interactions with *A. calidoustus*, we imaged wild-type *A. calidoustus* mycelium after co-incubation with *P. edwinii* strains in PDB using backscatter scanning electron microscopy (BSEM). Wild-type and complement strains of *P. edwinii* formed micro-aggregates of bacterial cells adhered to the fungal mycelium, while micro-aggregates were not observed in the Δ*fac* operon strain (Fig. 5A). Cells of the *fac* mutant strains were only observed sparsely and individually adhered to the fungal hyphae (Fig. 5B).

**Figure 5:**
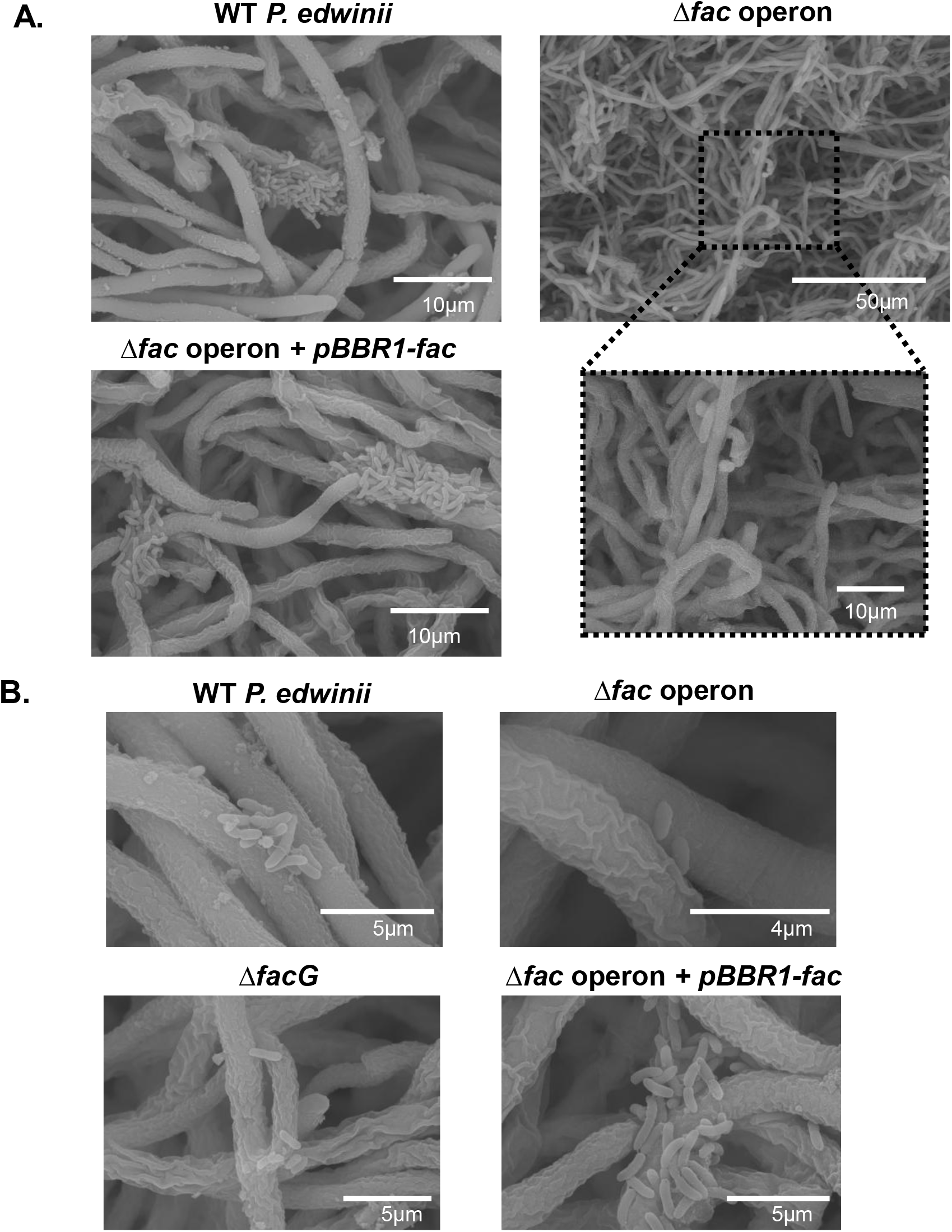
The *fac* system promotes micro-aggregate formation during co-incubation. Backscatter scanning electron microscopy (A) Wild-type and Δ*fac* operon + pBBR1-*fac* strains form micro-aggregates adhered to fungal mycelium after co-incubation. Scale bar is 10µm. Δfac operon does not form observed micro-aggregates. Scale bar is 50µm or 10µm. (B) Wild-type, Δ*fac* operon + pBBR1-*fac*, Δ*fac* operon, Δ*facG* strains adhered to fungal mycelium. Scale bar is 5 or 4 µm.

### The *fac* system is not essential for mature bacterial aggregate formation during the protection response

Since *P. edwinii fac* mutants had a PCA fungal protection deficit, consistent with the *fac* operon being under the negative regulatory influence of HrcA, we hypothesized the *fac* operon may play a role in the enhanced mature bacterial aggregation seen in the Δ*hrcA* strain (Fig. 1A), which was previously suggested to be associated with enhanced sequestration of PCA in this ‘toxin sponge’ bacterium (22). Additionally, the BSEM images suggest the *fac* system’s involvement in micro-aggregate formation upon initial fungal mycelium association (Fig. 5A-B), but it is unknown if this mechanism is shared with mature bacterial aggregate formation in the co-colony in the presence of PCA. To evaluate *fac* mutants for mature bacterial aggregation during the protection response, we developed an assay that allows visualization of the bacterial aggregates with the use of a stereoscope. The freshwater fish pathogen *Flavobacterium columnare* has been shown to increase aggregation formation on fish gills as a response to increasing concentrations of L-rhamnose (49), and L-rhamnose is available from plant-derived polysaccharides in rhizosphere environments (50), including environments similar to that from which *P. edwinii* was isolated. Thus, we tested if increased concentrations of L-rhamnose would enhance mature bacterial aggregation in co-colony in the presence of PCA. We found 5mM L-rhamnose produces mature bacterial aggregates clearly visible by stereoscope. Both the Δ*fac* operon and the Δ*facG* strains were able to produce bacterial aggregates comparable to wild-type under these conditions at 48 hours, indicating the *fac* system is not essential for the formation of mature bacterial aggregates during the protection response (Fig. 6A). These results also indicate the Δ*hrcA* hyper-aggregation phenotype is unlikely to be explained by the upregulation of the *fac* system, and these archaic CUP genes are primarily involved in adherence to host cells, and not adherence to self in the mature bacterial aggregates (29, 36).

**Figure 6:**
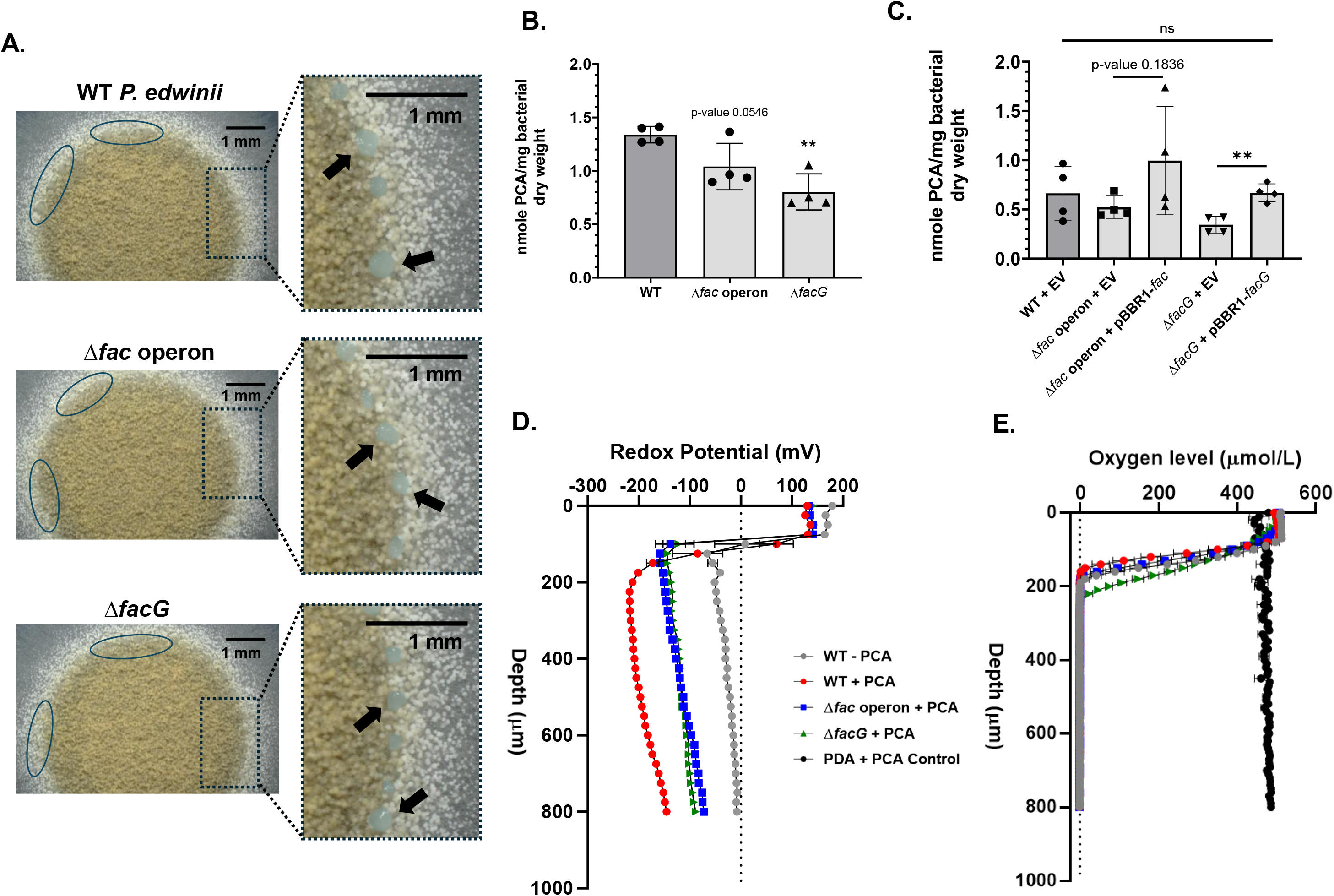
The *fac* system is involved in *P. edwinii’s* PCA sequestration and detoxification mechanisms. (A) Wild-type, Δ*fac* operon, Δ*facG* bacterial aggregation assay in co-colony on PDA with 300µM PCA and 5mM L-rhamnose at 48 hrs. Blue circles and black arrows indicate mature protection response bacterial aggregates. Cropped zoom-in’s bacterial aggregates are overlayed with a blue false color for visualization. Scale bar is 1 mm. Represents 3 biological replicates. (B-C) nmol of PCA sequestrated/mg of bacterial dry weight of wild-type, Δ*fac* operon, Δ*facG* strains, complement, and EV strains at 72 hrs in protection response activating conditions. Error bar is standard deviation. Represents four biological replicates. PCA quantified by UHPLC/MS. **p value<0.01. (C) One-way ANOVA with Dunnett’s posttest produced a non-significant result, while an unpaired t-test with Welch’s correction for unequal variance was used to compare mutant complementation to mutant + EV strains (D) Redox potential (mV) through the depth of mutant bacterial colonies in protection response activating conditions at 90 hrs. Measurements were taken every 25µm with three technical measurements. Represents two biological replicates. (E) Oxygen level (µmol/L) through the depth of mutant bacterial colonies in protection response activating conditions at 90 hrs. Measurements were taken every 10µm with three technical measurements. Represents two biological replicates.

### The *fac* system is involved in *P. edwinii’s* PCA sequestration and detoxification mechanisms

The final two characteristics of the Δ*hrcA* strain include an enhanced ability to sequester and detoxify PCA compared to wild-type (22). *P. edwinii* sequesters PCA and lowers the reduction potential of the local environment which may help slow the redox cycling of this phenazine and thus lower the level of ROS produced as a result (22). To understand if the *fac* system influences *P. edwinii’s* PCA sequestration efficacy, we quantified PCA sequestered by bacterial colonies in protection response activating conditions using ultra-high performance liquid chromatography mass spectrometry (UHPLC/MS). Both the Δ*fac* operon and the Δ*facG* strains are less effective at sequestering PCA than the wild-type strain (Fig. 6B). We complemented Δ*facG* from a plasmid, and it sequestered comparable amounts of PCA as wild-type, while the Δ*fac* operon showed partial complementation (Fig. 6C). These results indicate the *fac* system plays a role in PCA sequestration. Next, we evaluated the efficacy of *fac* mutants to detoxify PCA by measuring the bacterial colony’s internal redox potential in protection response activating conditions. The wild-type strain reached a redox potential of −212 mV, while the Δ*fac* operon and the Δ*facG* strains were only able to reach a redox potential of −138 mV, indicating the *fac* deletion strains produce a less reducing microenvironment than wild-type and therefore are less able to detoxify PCA in addition to being less efficient at sequestering the molecule in the first place (Fig. 6D and Supplementary Fig. 6).

Canonical archaic CUP pili are implicated in biofilm formation leading to a hypoxic microenvironment. This hypothetically links the *fac* mutants increase in redox potential to an increase in oxygen penetration. However, it has been demonstrated that the *fac* system is not necessary for bacterial aggregation, leading us to hypothesize oxygen concentration is not responsible for the increase in redox potential within the *fac* mutants. To explore whether the *fac* system maintains a reducing microenvironment by physically excluding oxygen through biofilm formation or if the *fac* system plays a more direct role in PCA detoxification, we measured the oxygen level of *fac* mutant colony biofilms using the same protection-activating conditions as measuring the local redox potential. The *fac* mutant’s oxygen gradient was virtually identical to that of wild-type (Fig. 6E and Supplementary Fig. 7), indicating the level of oxygen is not what is contributing to the more oxidizing environment produced by the mutant strains. Given that the *fac* system does not appear to be necessary to form mature bacterial aggregates, nor does its absence alter oxygen penetration into the colony, it is more likely that this non-canonical archaic CUP system is more directly involved in the activation and/or maintenance of PCA sequestration and reduction mechanisms.

### An AraC-family transcription regulator, FacR, influences *fac* operon expression in the presence of PCA

The RNA-seq revealed an AraC-family transcription regulator, *facR*, was also upregulated in the Δ*hrcA* strain (Fig. 1B). AraC-like transcription regulators have been implicated in the regulation of CUP operons, and *P. aeruginosa* CUP operon expression is influenced by regulatory cascades (43, 51, 52). Thus, we hypothesized *facR* may influence the expression of the *fac* system. The *facR* deletion strain had a PCA fungal protection defect when compared to the wild-type strain and could be complemented from a plasmid (Fig. 7A-B). We evaluated the Δ*facR* strain’s protection mechanisms: bacterial aggregation, PCA sequestration, and PCA detoxification. Similarly to the *fac* system mutants, the Δ*facR* strain was able to make bacterial aggregates on 5mM L-rhamnose but sequestered less PCA and was less efficient at detoxifying PCA compared to wild-type (Fig. 7C-E and Supplementary Fig. 7-8). Considering Δ*facR* had protection response defects similar to the Δ*fac* operon strain, we measured the *fac* operon expression level in the Δ*facR* background using quantitative real-time PCR (qRT-PCR). *fac* operon expression significantly decreased by ∼40% in the Δ*facR* strain compared to wild-type in the presence of PCA, indicating FacR influences the activation of *fac* operon expression. Significantly, *fac* operon expression level is unaffected by the deletion of *facR* in the absence of PCA, suggesting PCA influences the activation of FacR (Fig. 8A and Supplemental Fig. 8). To validate the qRT-PCR results, we evaluated if expression of the *fac* operon expressed from a plasmid in the Δ*facR* background could restore PCA fungal protection. The Δ*facR* + *pBBR1-fac* operon strain had increased fungal protection compared to the Δ*facR* + empty vector (EV) and wild-type + EV strains, compatible with a model where FacR influences the activation of the *fac* operon in the presence of PCA (Fig. 8B). Taken together, these results suggest that derepression of HrcA by PCA leads to upregulation of FacR, which in turn upregulates the *fac* operon that encodes components of an interkingdom protection system needed to protect a fungal partner, but not *P. edwinii* itself, from PCA (Fig. 8C).

**Figure 7:**
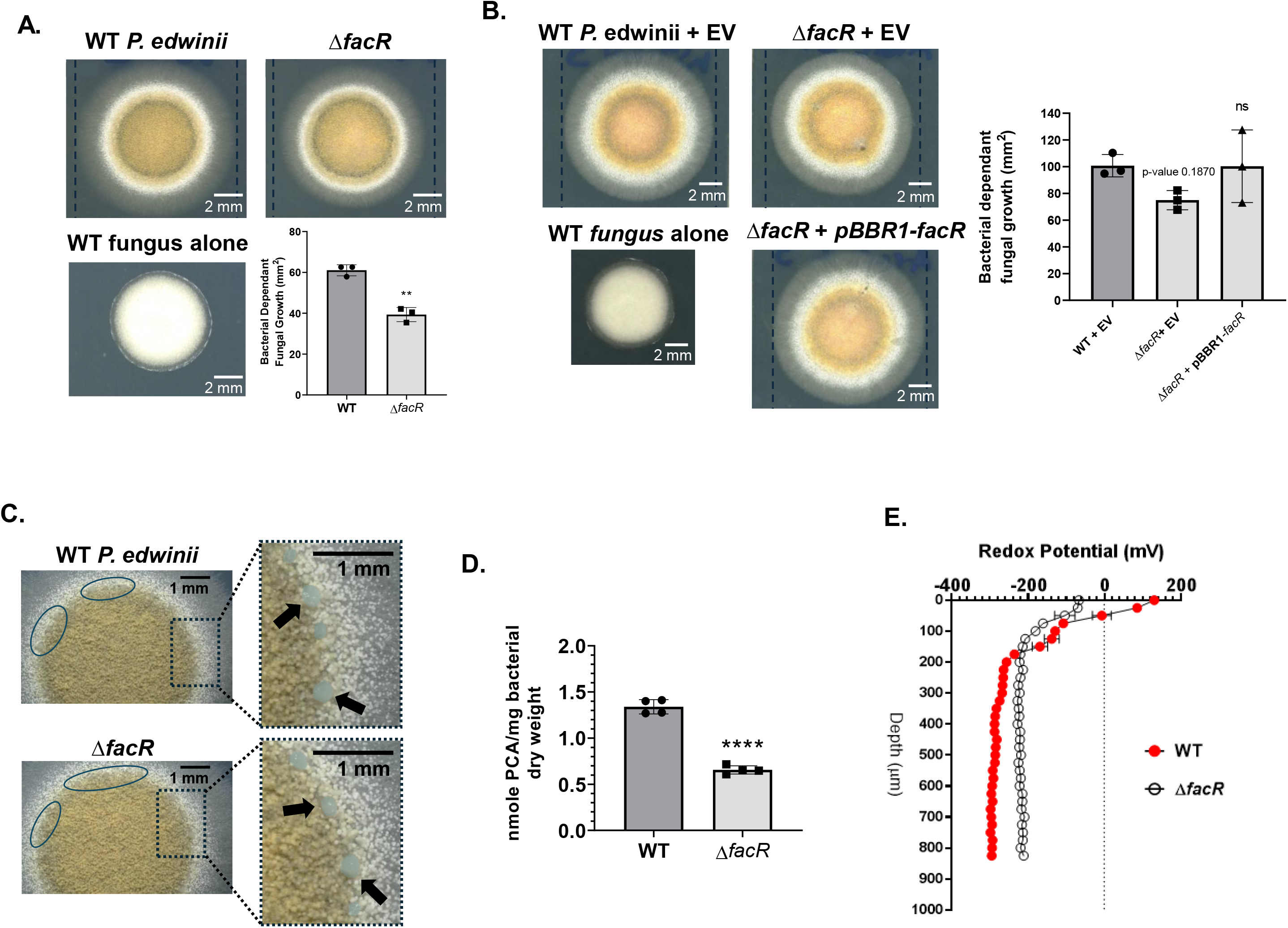
An AraC-family transcription regulator, FacR, plays a role in the bacterial protection response. (A-B) Since plasmid carriage impacts *P. edwinii* growth under these conditions, deletion mutants were compared to plasmid-free wild-type at 68 hrs, whereas complemented strains were compared to the corresponding wild-type + EV control at 90 hrs. Dashed lines indicate diameter of wild-type or wild-type EV co-colony. Represents three biological replicates. Scale bar indicates 2 mm, and error bars indicate standard deviation. Y-axis indicates bacterial-dependent fungal growth, with zero indicating the average area of three biological replicates of *A. calidoustus* grown alone on PCA. *p value<0.05, **p value<0.01. (A) *facR* mutant in co-colony displays a statically significant decrease in bacterial dependent fungal growth on 300µM PCA compared to wild-type. (B) *facR* complement strain in co-colony grown on PDA with 300µM PCA. Gene expression from pBBR1 under a rhamnose inducible promoter. (C) Δ*facR* bacterial aggregation assay in co-colony on PDA with 300µM PCA and 5mM L-rhamnose at 48 hrs. Blue circles and black arrows indicate mature protection response bacterial aggregates. Cropped zoom-in’s bacterial aggregates are overlayed with a blue false color for visualization. Scale bar is 1 mm. Represents 3 biological replicates. (D) nmol of PCA sequestrated/mg of bacterial dry weight of Δ*facR* at 72 hrs in protection response activating conditions. Error bar is standard deviation. Represents four biological replicates. PCA quantified by UHPLC/MS. ****p value<0.0001. (E) Redox potential (mV) through the depth of mutant bacterial colonies in protection response activating conditions at 90 hrs. Measurements were taken every 25µm with three technical measurements. Represents two biological replicates.

**Figure 8:**
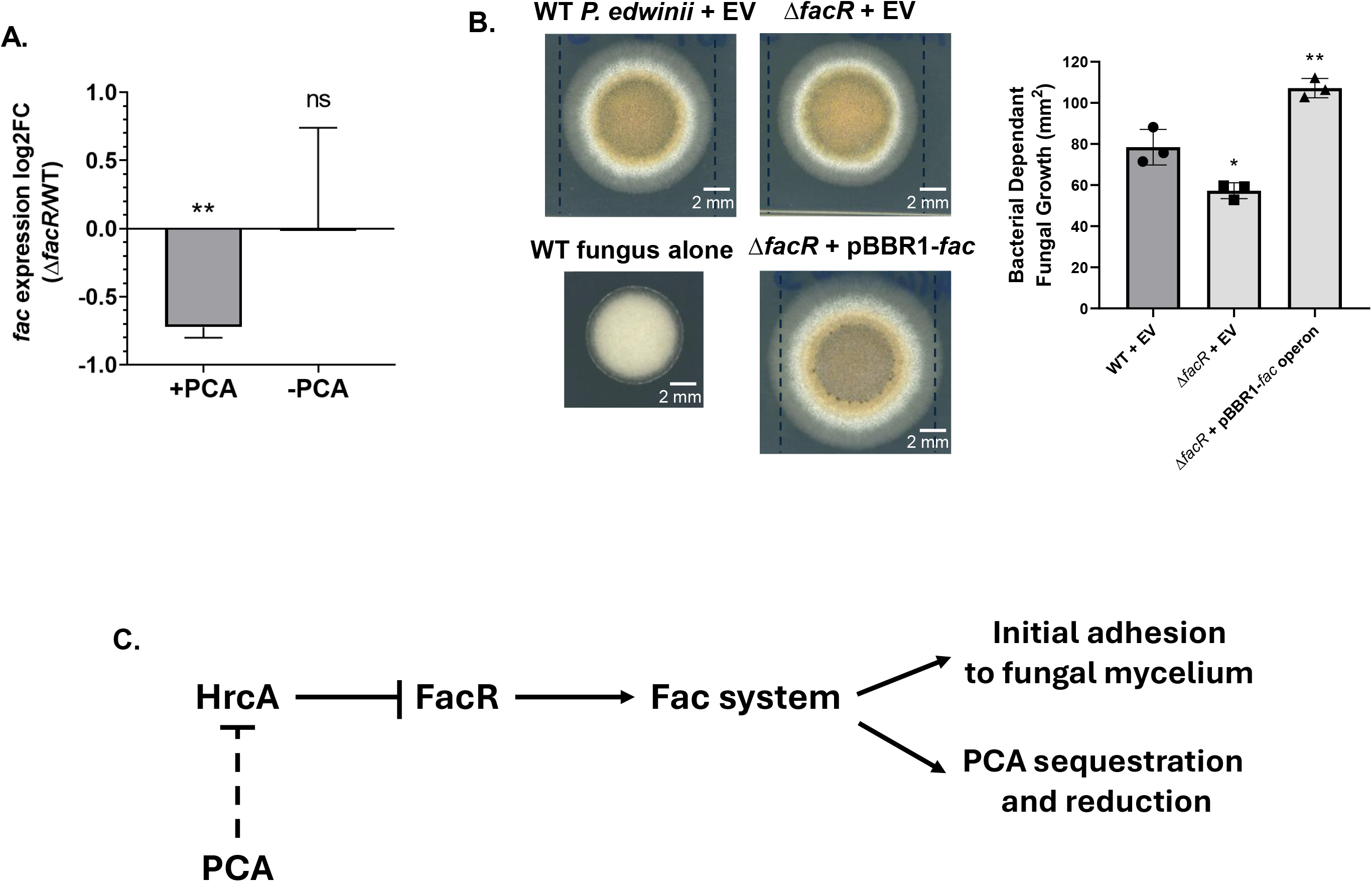
FacR influences the expression of *fac* system in the presence of PCA. (A) *fac* operon expression, measured by qRT-PCR, is significantly decreased in the Δ*facR* background in the presence of PCA at 12 hrs compared to wild-type. **p value<0.01. (B) *fac* operon expressed from a rhamnose inducible promote in the Δ*facR* background in co-colony on 300µM PCA and 100µM L-rhamnose restored bacterial dependent fungal growth. Y-axis indicates bacterial-dependent fungal growth, with zero indicating the average area of three biological replicates of *A. calidoustus* grown alone on PCA. *p value<0.05 and **p value<0.01. (C) Model of purposed *fac* system regulatory pathway. PCA is thought to inhibit HrcA directly or indirectly to activate the protection response, which represses FacR. FacR activates the expression of the *fac* system in the presence of PCA. The *fac* system is involved in bacterial-fungal association and PCA sequestration and reduction.

## DISCUSSION

Microbial communities often contain organisms that are sensitive to antimicrobial compounds produced by their neighbors, suggesting that community membership can depend on protective interactions as well as intrinsic tolerance (19, 20). Here, we used the recently developed model system in which the bacterium *P. edwinii* acts as a toxin sponge to protect its physically associated fungal partner *A. calidoustus* from PCA to determine if genetic elements responsible for this response are involved in self-protection or specifically in interkingdom-protection. We used an *hrcA* deletion strain where the protection response is locked in the active state to determine which genetic elements in *P. edwinii* might be responsible for HrcA’s apparent control over four characteristics of fungal protection: adhering to fungal cells, formation of mature bacterial aggregates within the fungal mycelium, PCA sequestration, and PCA detoxification by reduction (22).

We show that a putative archaic CUP system we termed **f**ungal **a**ttachment **c**omplex (*fac*) appears to play a role in three of these four protection characteristics during phenazine stress. Non-canonical archaic CUP operons are widespread in the bacterial domain, and we currently have a limited understanding of how closely their functions mirror canonical CUP pili, spore coat, or capsule functions (24). In this study, we demonstrate that a non-canonical archaic CUP system is involved in the bacterial protection of its fungal partner from AMCs. This helps expand known adhesion targets from animal host or abiotic surfaces to fungal cells, and broadens the function of such systems from being involved in self-protection to AMCs through biofilm formation or environmental stress via spore coat to interkingdom protection through association with a eukaryotic partner and involvement in detoxification of AMCs (36, 37).

Archaic CUP pili have been primarily studied in the human pathogens *A. baumannii* and *P. aeruginosa*, where they are primarily thought of as virulence factors contributing to abiotic adhesion, host cell adhesion, and self-protecting biofilms that promote antimicrobial resistance (29, 35, 36). However, the archaic clade is also considered the most ancestral and diverse, containing operons that encode secreted pili, non-pili secreted factors such as the *M. xanthus* spore-coat protein, and many divergent operons whose function are as of yet undetermined (24, 26). These previous sets of observations made the *fac* system a compelling target for study as there was reason to think that pili or adhesive structures could be involved in all four regulated characteristics of *P. edwinii*’s protection response. Intriguingly, we did not find evidence that the *fac* system is relevant for self-adherence in the formation of mature bacterial aggregates in the fungal mycelium. Mature aggregate formation may instead involve alternative adhesion systems, such as MipA/OmpV, YadA-like family protein, or OmpA family protein which were also significantly upregulated in the Δ*hrcA* at 12hr but did not meet our log2FC > 1 cutoff (53–55). Instead, the *fac* system is important for adherence to fungal cells upon initial association, promotion of PCA sequestration during the ‘toxin sponge’ response, and plays a role in producing or maintaining the reducing environment that allows *P. edwinii* to detoxify PCA. Even more surprising, the *fac* system is not required for *P. edwinii* self-protection from PCA stress, thus demonstrating that archaic CUP systems can have a broader range of functions that encompass more than self-protection via biofilm or spore-coat formation, and may have functions that extend past physical adhesion and cohesion.

Adhesion to the fungal partner is the first step in protection, and this study demonstrates the *fac* system is necessary for adhesion to fungal cells and initial bacterial-fungal association. To our knowledge, this is the first evidence that an archaic CUP system promotes bacterial adherence to fungal cells, broadening the known functional scope of the archaic clade. The fact that the *fac* system is not required for mature bacterial aggregation in co-culture, but is required for adherence to fungal mycelium may help answer another important question. Previous work suggests that while protective bacteria show modest discrimination about which fungi they will protect from AMCs, the biggest selective step is association with a given fungal host in the first place (22, 56). The results show the *fac* system provides fast adherence to fungal mycelium provides an important first hypothesis for the mechanism of how association occurs in nature. *Paraburkholderia unamae* has been shown to protect *A. calidoustus* from PCA stress despite not being found associated with *A. calidoustus* in nature (22). Interestingly, *P. unamae* (taxid: 219649) does not contain a homologue of the *fac* system in its genome, suggesting that natively associated partnerships may have developed specialized extracellular mechanisms for protection. Providing further evidence the *fac* system shares adherence functions with pili, the BSEM images in this study show the wild-type and complement strains of *P. edwinii* associating with *A. calidoustus* hyphae as micro-aggregates, while the *fac* mutants failed to form micro-aggregates and instead were seen sparsely and individually adhered to the fungal hyphae. The function of micro-aggregate formation upon initial association may be to allow for quick and robust colonization of the fungal partner in moist and turbid environments. However, if these micro-aggregates form before or after adhesion to the fungal mycelium has yet to be determined, and some studies indicate archaic CUP pili cause pellicle formation in liquid media, which may explain the patchwork initial adherence of wild-type *P. edwinii* (36). Interestingly, filamentous structures are seen spanning the micro-aggregates that are suggestive of bundles of antiparallel stacked archaic pili (36) (Fig. 4B). However, it is plausible the observed filamentous structures are an artifact of SEM, therefore we explored imaging a singular bacterium from monoculture for a pilus structure using TEM. However archaic CUP pili are notoriously ultrathin structures (2.3 to 6 nm) that are difficult to resolve (34, 35). Visualization of a pilus structure is further complicated by the presence of multiple annotated fimbriae loci, making it difficult to attribute observed surface appendages directly to the *fac* system.

After initial adherence to fungal mycelium and the formation of mature bacterial aggregates in co-colony, the final steps involved in protection of *A. calidoustus* from PCA are sequestration and detoxification of this molecule. While this study implicates the *fac* system in these two processes, the mechanism by which the *fac* system is able to help sequester PCA and maintain a low reducing potential microenvironment to detoxify it remains to be fully elucidated. Archaic CUP pili have been heavily implicated in biofilm formation in human pathogens, and biofilms have decreasing oxygen concentrations (29, 36, 37), and therefore one hypothesis is that the *fac* system is acting to promote mature bacterial aggregation and thus contribute to the creation of the anoxic, highly reduced zone necessary for PCA detoxification. However, the *fac* system was not required for mature bacterial aggregate formation, which may be driven by one of the six other pilus/fimbriae systems annotated in *P. edwinii’s* genome, and PCA sequestration efficacy was disrupted despite functional mature bacterial aggregation. Additionally, the *fac* mutants did not differ from wild-type in terms of oxygen depletion in a colony biofilm, despite producing a defect in the local reduction potential under the same protection activating conditions (Fig. 5). This decoupling effect suggests the *fac* system is involved in contributing to the protection program in a more direct manner. Considering type I and type IV pili have been described acting as mechano-sensors (30, 57, 58) one hypothesis is that archaic CUP systems may also be involved in chemical or mechanical signaling processes. An alternative hypothesis is that the *fac* system plays a structural role in mature bacterial aggregates, helping to retain sequestered PCA and/or to limit the diffusion of locally produced metabolites, such as organic acids, that contribute to reduction of the surrounding microenvironment.

A final question this study raises is if the *fac* system behaves in a manner that is consistent with it being a pilus as opposed to an alternative extracellular product. This study provides evidence that the *fac* system shares features with pili, including roles in AMC resistance and host cell adherence, as well as the predicted structure of the putative adhesive tip, FacG, includes a hydrophobic pocket, a feature of known pilus adhesive tips (44). However, critical differences remain including the *fac* system aiding the host to resist AMCs instead of its bacterial parent cell, and aiding in the sequestration and detoxification of an AMC. Additionally, the protein structure secreted by the *fac* system has yet to be resolved, and therefore it is possible that this operon encodes for a distinct class of secreted protein similar to the F1 capsule secreted by *caf1* system in *Y. pestis* (25) or the spore-coat protein secreted by MXAN_3885-2 system in *M. xanthus* which shares the COG5430 conserved domain with FacA and FacG subunits (26, 43). The fact that the *fac* system shares features with the MXAN_3885-2 system and participates in unique functions reveals future biochemical, structural, and localization studies should focus on determining if the *fac* system secretes a pilus or an alternative extracellular product. Additional work is also necessary to fully understand how *P. edwinii* forms large mature aggregates in the fungal mycelium, identifying the specific substrate of FacG’s hydrophobic binding pocket, and determining the if *fac* system act as a sensor to promote a low local reduction potential and any downstream targets in a possible signaling cascade.

Finally, we identified an AraC-family transcriptional regulator, FacR, that influences the activation of the *fac* system in the presence of PCA. Previous work has shown that HrcA is likely a negative repressor of the protection response, and derepression during PCA exposure is partially responsible for activation of the response (22). Given that components of the *fac* system and *facR* were both upregulated in the Δ*hrcA* background, and FacR upregulates the *fac* system in the presence of PCA but not in the absence of PCA, suggests HrcA, FacR, and the *fac* system participate in the same protection response pathway (Fig. 1B & 6C). AraC-family transcription factors have been well documented to influence CUP expression in various microbes and transcription regulatory cascades have been reported in *P. aeruginosa*, making it plausible for the *fac* system to be regulated by a transcription factor cascade (43, 52). We therefore view the likely cascade as PCA deactivating HrcA repression, leading to FacR upregulation, which in turn upregulates the distal *fac* system operon. That adding PCA can cause an increase in transcription of the *fac* operon in wild-type, but not in a *facR* mutant, is strong evidence in favor of this view.

The identification of an archaic CUP system involvement in protective microbe-microbe relationships highlights the functions of archaic CUP operons beyond self-action or as virulence factors, revealing the diverse functional niches archaic CUP systems may fill. The *fac* system is not necessary for self-protection of the bacteria that produce them in this system but instead extends interkingdom protection to a fungal partner, while retaining overlapping functions with known pili. Given diverse protective bacterial-fungal relationships have been identified over recent years (22, 56), this mechanism may be prevalent in agricultural and infection site contexts. This in turn may aid in further deciphering how microbial communities that drive health or pathogenesis in plants and humans form in a species-specific manner, informing agricultural practices and clinical drug choices. Finally, elucidating how the *fac* system is used during initial protective bacterial-fungal association and protection response activation will allow us to manipulate beneficial fungal AMC tolerance, or pathogenic fungal AMC susceptibility, through their bacterial partners, thus opening new pathways to treat recalcitrant infections and deal with food insecurity by tailoring a crop’s rhizosphere to changing climate conditions. Understanding the mechanisms that are involved in microbial community membership will allow us to view critical communities through a more dynamic lens that is reflective of the species-level composition of those communities that define their impacts on plants and people.

## METHODS

### Construction of bacterial strains and plasmids

In-frame deletion strains were constructed using cloning tools adapted from *Pseudomonas* spp. (59). Constructed pmQ30 suicide plasmids were transformed by electroporation into *E. coli* S17 strains, then conjugated into wild-type *P. edwinii* on yeast peptone agar plates (YPA) overnight at 30C. Mated cells were grown on PDA containing gentamicin and chloramphenicol for three days at 30C to select for *P. edwinii* containing pmQ30. Strains were then grown on YPA plates containing 7.5% sucrose for two days at 30C to select from cells that underwent homologous recombination. Mutant strains were PCR confirmed and mutants in main figures were amplicon sequence confirmed. Expression plasmids were constructed similarly, and a rhamnose inducible promoter system was inserted into the broad-spectrum plasmid, pBBR1. Correctly constructed expression plasmids were sequenced confirmed and electroporated into *P. edwinii* strains and selected for on kanamycin. Details are in the Supplemental Materials and Methods.

### Growth conditions

All *P. edwinii* or *A. calidoustus* cultures were grown in 5ml PDB or PDA plates at 30C. Liquid cultures were rotated at 70rpm and 150µM of PCA and/or L-rhamnose was added when indicated. PDA plates contained 300µM PCA and 250µM for complementation or 5mM L-rhamnose when indicated. Protection activating conditions indicate *P. edwinii* strains grown 5mm from wild-type *A. calidoustus* on PDA containing 300µM PCA. Growth time point is indicated in specific assay method section and figure legends.

### RNA-sequencing

WT and Δ*hrcA* strains of *P. edwinii* were normalized to 0.2 OD600, plated on PDA, and grown at 30C for 6, 12, and 24 hours in three biological replicates. Colonies were then scraped up with PDB and pelleted. RNA was extracted and DNase treated. RNA quality was evaluated by Qubit, electrophoresis, and Agilent Bioanalyzer. High quality RNA was ribodepleted, library prepped, pooled, then sequenced using NextSeq 2000 (paired end, 100 cycles, 400M). RNA-seq analysis was conducted using the Galaxy EU biological analysis, HISAT2, FeaturedCounts, and DESeq2 (60). Volcano plots were constructed in R-studio. Details are in the Supplemental Materials and Methods.

### Protein predictions and cladogram construction

SignalP was used to identify N-terminal signal peptide sequences that would be cleaved in mature protein (61). Google AlphaFold3 server was used to predict the mature protein structure and ChimeraX was used to visualize the CIF output files. CUP outer membrane usher amino acid sequences with known classification were gathered using NCBI data base and Nuccio and Baumler, 2007 (24). Amino acid alignment and cladogram construction was done using MEGA12 software using maximum likelihood and adaptive bootstrap analysis.

### Co-culture protection assay

*P. edwinii* strains were normalized to 2.5×10^-4^ OD600. Bacterial cells were mixed 1:1 with 25 million spores/mL of wild-type *A. calidoustus* spores prep. Wild-type *A. calidoustus* spores were also mixed 1:1 with 1x sterile PBS to serve as a negative monoculture control. Six µl of each co-culture or control were plated on PDA containing 300µM of PCA, and 250µM L-rhamnose for complementation strains. At 68 or 90 hours of growth co-colonies were imaged using the Keyence VHX digital microscope. Fungal growth was quantified by measuring co-colony areas using the Keyence area analysis tool. The average area of three biological replicates of *A. calidoustus* alone were on PCA were subtracted from the co-colony area of three biological replicates to produce the area of bacterial-dependent fungal growth. Error bars indicate standard deviation and a One-way ANOVA with Dunnett’s posttest was conducted for multiple comparisons or an unpaired t-test with Welch’s correction for unequal variance for a single comparison to calculate statistical significance between samples (p-value).

### Bacterial CFU quantification

Co-cultures and bacterial monocultures were prepared the same as above and grown on PDA with and without 300µM PCA for 48 hours. Colonies were cut out of the agar put into 4ml of Yeast Peptone (YP) broth. Colonies were homogenized for 1 minute at 60% amplitude. Homogenized samples were diluted 10^-1^ to 10^-5^ in 100µl of YP. Ten µl of each co-culture or monoculture dilution were spotted at the top of YP plates with or without 50µg/ml of Nystatin and dripped down plate. Drip plates were placed in 30C for 48hr. Bacterial CFUs were counted and the total number of bacterial CFUs in the co-culture or monoculture spots were back-calculated. Total bacterial CFUs were averaged between four biological replicates. Error bars indicate standard deviation and a One-way ANOVA with Dunnett’s posttest was conducted for multiple comparisons or an unpaired t-test with Welch’s correction for unequal variance for a single comparison to calculate statistical significance between samples (p-value). Details are in the Supplemental Materials and Methods.

### Redox potential and oxygen concentration probes

Monocultures of the bacterial strains and wild-type *A. calidoustus* were prepared the same as above, but plated 5mm apart (side by side) on PDA containing 300µM PCA to induce the protection response and grown at 30C. At 90hr of growth the bacterial colonies were probed using the Unisense redox microelectrode paired with a reference probe and the oxygen microelectrode. Three technical redox measurements were taken at every 25µm for a total of 1000µm, and three technical oxygen measurements were taken every 10µm for a total of 1000µm into the bacterial colony of two biological replicates. Details are in the Supplemental Materials and Methods.

### Ultra-High Performance Liquid Chromatography Mass Spectrometry (UHPLC/MS)

*P. edwinii* and *A. calidoustus* monocultures were prepared, plated side by side, and grown the same as above. At 68hr of growth the bacterial colonies were scraped up and pelleted. Bacterial pellets were resuspended in 200µl of 1x sterile PBS and pelleted again. The bacterial pellets were placed in −80C before overnight lyophilization. The supernatant was moved into a fresh tube and placed in the −20C overnight to allow for larger biotic particles to sediment. After a freeze thaw cycle, 150µl of the supernatant was transferred to a 0.22µm cellulose acetate Spin X column. The column was centrifuged at 13,000xg for 1 minute and the flow-through of four biological replicates was sent for liquid chromatography mass spectrometry (UHPLC/MS) at the University of Georgias Proteomics and Mass Spectrometry facility. Samples were analyzed on a Bruker Impact II mass spectrometer interfaced with a Bruker Elute UHPLC. The UHPLC was equipped with a Waters XBridge Premier BEH C18, 3.5 µm, 21. x 50 mm column. Three technical measurements were taken, and the data was analyzed using Bruker Compass Data Analysis. PCA quantification was normalized to bacterial dry weight. Error bars indicate standard deviation and a One-way ANOVA with Dunnett’s posttest was conducted for multiple comparisons or an unpaired t-test with Welch’s correction for unequal variance for a single comparison to calculate statistical significance between samples (p-value). Details are in the Supplemental Materials and Methods.

### Aggregation Assay

Co-cultures were prepared the same way as the protection assay, except *P. edwinii* strains are normalized to 2.5×10^-3^ OD600. Co-cultures were plated on PDA containing 300µM PCA and 5mM L-rhamnose for 48hr at 30C. Three biological replicates were imaged for mature bacterial aggregation using the Keyence VHX digital microscope.

### Adhesion Assay

1.25 million wild-type *A. calidoustus* spores were grown in 5ml of PDB overnight for approximately 12 hrs. *P. edwinii* strains were normalized to 0.25 OD600 and 50µl were inoculated into the 5ml of PDB containing young wild-type *A. calidoustus* mycelium growth. *P. edwinii* and *A. calidoustus* were incubated together for 30 minutes or 3 hours at 30C, 70rpm. Mycelium was washed with 5ml of PDB twice and transferred to 500µl of fresh PDB. Mycelium was homogenized for 1 minute at 65% amplitude. Homogenized samples were diluted 10^0^ to 10^-4^ in 100µl of YP. Ten µls of each dilution were spotted at the top of YP plates containing 50µg/ml of Nystatin and dripped down. Drip plates were placed in 30C for 48hr. Bacterial CFUs were quantified the same as above. Total number of bacterial CFUs were averaged between four to six biological replicates. Error bars indicate standard deviation and a One-way ANOVA with Dunnett’s posttest was conducted for multiple comparisons or an unpaired t-test with Welch’s correction for unequal variance for a single comparison to calculate statistical significance between samples (p-value). Details are in the Supplemental Materials and Methods.

### Backscatter Scanning Electron Microscopy (BSEM)

*P. edwinii* and *A. calidoustus* co-incubation cultures was prepared the same way as the adhesion assay and co-incubated for 30mins to 1.5hr at 30C, 70rpm. *A. calidoustus* mycelium was washed with 5ml of 1x sterile PBS twice, transferred to 500µl of 2% paraformaldehyde in 1x PBS, and sent to Georgia Electron Microscopy facility for imaging. Cell suspensions were affixed to coverslips for BSEM imaging with a protocol adapted from Mazia et al.(1975) (62). Samples were sputter-coated with a 30 nm layer of gold-palladium and viewed on a Hitachi SU3900 SEM. Details are in the Supplemental Materials and Methods.

### Quantitative reverse-transcriptase PCR (qRT-PCR)

RNA from three biological replicates was prepared in the same way as the RNA-seq samples, with samples normalized to total RNA. Reverse transcriptase (RT) and no RT reactions were conducted using SuperScript IV VILO Master Mix. qPCR was conducted on each sample’s cDNA with primers designed to target the transcript of interest. Amplification of RpoD was used as a stable endogenous control. Target primer efficiency was confirmed by standard curve using known concentrations of cDNA. The Azure Cielo qPCR systems quantified each reaction’s CT value. CT values are converted to log2FC using Livak and Schmittgen method (63). The Log2FC average and standard deviation was calculated of three biological replicates. An unpaired t-test with Welch’s correction for unequal variance was conducted to calculate statistical significance between samples (p-value).

## Data availability

The data from this study is available on request from the corresponding author. RNA-seq data is available on NCBI SRA (PRJNA1516646). AlphaFold structure prediction data and UHPLC/MS data will be submitted to Zenodo upon acceptance.

## Acknowledgements

We would like to thank the Dahlstrom lab members for project feedback, specifically Hannah Edwards and Renuka Namala. We thank Rochelle Yap and Abigail Deaven, for RNA-seq analysis guidance, Alejandro Pereira and Logan Lafave, for structure analysis guidance. We would like to thank my dissertation committee, Courtney Ellison, Zack Lewis, and Mary Ann Moran. Jorge Escalante and Chelsey VanDrisse for reviewing the manuscript draft. RNA sequencing was done Georgia Genomics and Bioinformatics core. Electron microscopy was done at Georgia Electron Microscopy core by Bayleigh Roussel. UHPLC/MS was done at the Georgia Proteomics and Mass Spectrometry facility by Dennis Philips. This project was funded by the National Institute of Health (NIH) MIRA award: 1 R35GM150797-01 and the NIH T32 Genetics Training Grant:1T32GM142623. The content is solely the responsibility of the authors and does not necessarily represent the official views of the NIH.

## Author contributions

Alyssia Gonzalez conducted the experiments and wrote the original draft. Kurt Dahlstrom supervised the project and provided the initial research question. Alyssia and Kurt developed assays, came up with a research strategy, analyzed data, acquired funding for the project, re-wrote, reviewed and edited the paper.

