## Supplemental figures for "A bacterial chaperone–usher pathway links interkingdom association to antimicrobial protection of a fungal partner"

### Supplemental Figures for Fac System Manuscript (High-Quality Images)

Alyssia Gonzalez

September 2026

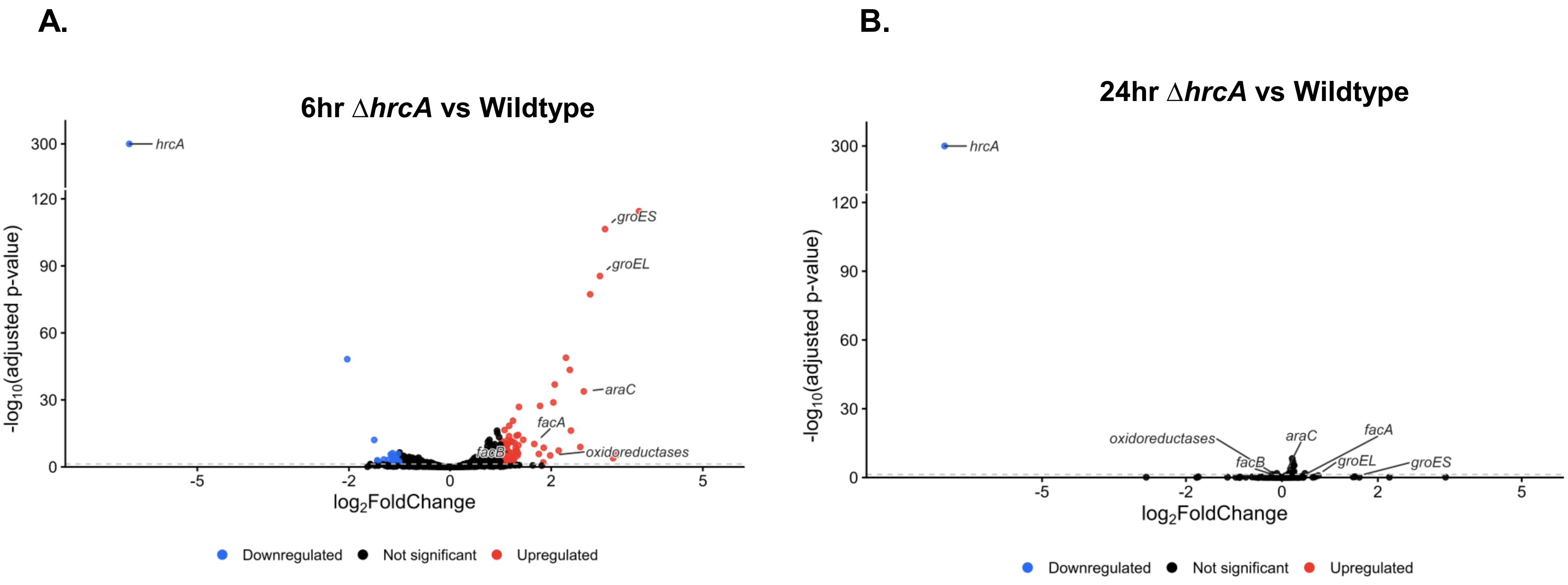


**Supplemental Figure 1:**
Differential gene expression in Δ*hrcA P. edwinii* strain compared to wild-type at (A) 6 and (B) 24 hours on PDA.

**Supplemental Table 1:**
Forty most significantly upregulated transcripts in Δ*hrcA P. edwinii* strain compared to wild-type at 12 hours on PDA. Remaining 69 significantly upregulated transcripts found in corresponding excel file.

| GeneID | GeneName | BaseMean | log2FoldChange | StdErr | WaldStats | PValue | PAdj |
|---|---|---|---|---|---|---|---|
| KZJ38_16360 | groES | 49031.41023 | 3.585192888 | 0.164904411 | 21.74103695 | 8.398E-105 | 3E-101 |
| KZJ38_16355 | groL | 80585.48074 | 3.508663452 | 0.168188629 | 20.86147843 | 1.19907E-96 | 2.36E-93 |
| KZJ38_14505 | entericidin A/B family lipoprotein | 1997.730024 | 3.506790742 | 0.46466938 | 7.546851365 | 4.45907E-14 | 7.38E-12 |
| KZJ38_15670 | alpha/beta hydrolase | 1302.979705 | 3.451200568 | 0.17579124 | 19.63238078 | 8.17909E-86 | 9.7E-83 |
| KZJ38_34150 | BON domain-containing protein | 735.04843 | 2.863848026 | 0.59816309 | 4.787737781 | 1.68672E-06 | 9.23E-05 |
| KZJ38_15650 | alpha/beta fold hydrolase | 363.2775153 | 2.75538104 | 0.155615647 | 17.70632381 | 3.74764E-70 | 3.33E-67 |
| KZJ38_16350 | hydroxymethylpyrimidine/phosphomethylpyrimidine kinase | 4467.059478 | 2.747641044 | 0.131739158 | 20.85667688 | 1.32569E-96 | 2.36E-93 |
| KZJ38_12040 | NADP-dependent oxidoreductase | 224.7955824 | 2.61925512 | 0.22791892 | 11.49204777 | 1.44642E-30 | 8.58E-28 |
| KZJ38_12245 | cytochrome P460 family protein | 4751.624127 | 2.592059688 | 0.163220327 | 15.88074071 | 8.6155E-57 | 6.81E-54 |
| KZJ38_33260 | YSC84-related protein | 1117.804918 | 2.576127809 | 0.513933104 | 5.012574183 | 5.37067E-07 | 3.18E-05 |
| KZJ38_15655 | AraC family transcriptional regulator | 134.6683807 | 2.558925532 | 0.228136315 | 11.21665147 | 3.37821E-29 | 1.85E-26 |
| KZJ38_23495 | NnrS family protein | 3321.840258 | 2.462996703 | 0.120724885 | 20.40173167 | 1.61398E-92 | 2.3E-89 |
| KZJ38_34145 | RNA polymerase factor sigma-54 | 1379.605697 | 2.436108332 | 0.251337152 | 9.692591442 | 3.24196E-22 | 9.61E-20 |
| KZJ38_22480 | EamA family transporter | 1737.205184 | 2.231775459 | 0.116883094 | 19.09408273 | 2.8278E-81 | 2.87E-78 |
| KZJ38_12290 | spore coat U domain-containing protein | 2851.770159 | 2.216703686 | 0.216177301 | 10.2541001 | 1.13428E-24 | 4.25E-22 |
| KZJ38_12045 | nitroreductase family protein | 366.1250758 | 2.21648722 | 0.214519305 | 10.33234384 | 5.03158E-25 | 1.99E-22 |
| KZJ38_14500 | DUF1328 domain-containing protein | 69.84822337 | 2.19159308 | 0.350245516 | 6.257305176 | 3.91686E-10 | 4.1E-08 |
| KZJ38_07760 | DUF3309 family protein | 107.5775995 | 2.154612563 | 0.326477249 | 6.599579502 | 4.12326E-11 | 4.97E-09 |
| KZJ38_08065 | hypothetical protein | 150.9594807 | 2.120142175 | 0.285488259 | 7.426372566 | 1.11616E-13 | 1.81E-11 |
| KZJ38_14510 | DUF4397 domain-containing protein | 194.969374 | 2.073379962 | 0.255863338 | 8.103466389 | 5.3415E-16 | 1.03E-13 |
| KZJ38_12050 | carboxymuconolactone decarboxylase family protein | 234.7245307 | 2.061678777 | 0.203963011 | 10.10810131 | 5.08604E-24 | 1.72E-21 |
| KZJ38_34160 | hypothetical protein | 224.0316819 | 2.021841082 | 0.288214986 | 7.015044955 | 2.29875E-12 | 3.48E-10 |
| KZJ38_32495 | autotransporter-associated beta strand repeat-containing protein | 3095.736358 | 1.966494348 | 0.452938739 | 4.341634265 | 1.41427E-05 | 0.000585 |
| KZJ38_23500 | AraC family transcriptional regulator | 1578.879646 | 1.888874573 | 0.177004715 | 10.67132349 | 1.38637E-26 | 6.17E-24 |
| KZJ38_34390 | helix-turn-helix domain-containing protein | 1731.162675 | 1.876660858 | 0.268574936 | 6.987475783 | 2.79876E-12 | 4.15E-10 |
| KZJ38_12295 | fimbria/pilus periplasmic chaperone | 446.7776435 | 1.79026485 | 0.188697675 | 9.487476999 | 2.36694E-21 | 6.74E-19 |
| KZJ38_18685 | DUF2282 domain-containing protein | 337.8403989 | 1.781936287 | 0.16760002 | 10.63207683 | 2.11353E-26 | 8.85E-24 |
| KZJ38_12035 | hypothetical protein | 269.2784542 | 1.767785333 | 0.173419787 | 10.19367721 | 2.11605E-24 | 7.53E-22 |
| KZJ38_11970 | cupin domain-containing protein | 415.681857 | 1.74209084 | 0.208614046 | 8.35078403 | 6.78113E-17 | 1.38E-14 |
| KZJ38_14490 | ferritin-like domain-containing protein | 408.4808201 | 1.691672535 | 0.290020793 | 5.832935339 | 5.44607E-09 | 4.78E-07 |
| KZJ38_12180 | cytochrome C oxidase subunit II | 81.29976104 | 1.655687564 | 0.315807202 | 5.24271629 | 1.5823E-07 | 1.06E-05 |
| KZJ38_18965 | hypothetical protien | 1529.047306 | 1.652678062 | 0.182577464 | 9.05192801 | 1.40466E-19 | 3.62E-17 |
| KZJ38_03200 | DUF883 family protein | 611.3807025 | 1.635010673 | 0.266492635 | 6.135294027 | 8.50018E-10 | 8.29E-08 |
| KZJ38_32095 | DUF1328 family protein | 40.70975954 | 1.608296778 | 0.474572007 | 3.388941519 | 0.00070163 | 0.017276 |
| KZJ38_10810 | DUF6723 family protein | 82.6601764 | 1.582876216 | 0.35058911 | 4.514904113 | 6.33454E-06 | 0.000304 |
| KZJ38_06990 | CsbD family protein | 4978.570833 | 1.581252739 | 0.19676887 | 8.036091989 | 9.27491E-16 | 1.69E-13 |
| KZJ38_21210 | rpoH | 36035.99485 | 1.561053525 | 0.115119124 | 13.56033187 | 6.8824E-42 | 4.9E-39 |
| KZJ38_11965 | cupin domain-containing protein | 43.67939737 | 1.560907535 | 0.356338235 | 4.380409906 | 1.18456E-05 | 0.000505 |
| KZJ38_34405 | helix-turn-helix domain-containing protein | 1162.60342 | 1.537935101 | 0.465070825 | 3.306883633 | 0.000943401 | 0.022083 |
| KZJ38_15235 | otsA | 4774.681774 | 1.526066494 | 0.246724862 | 6.185296775 | 6.19857E-10 | 6.3E-08 |

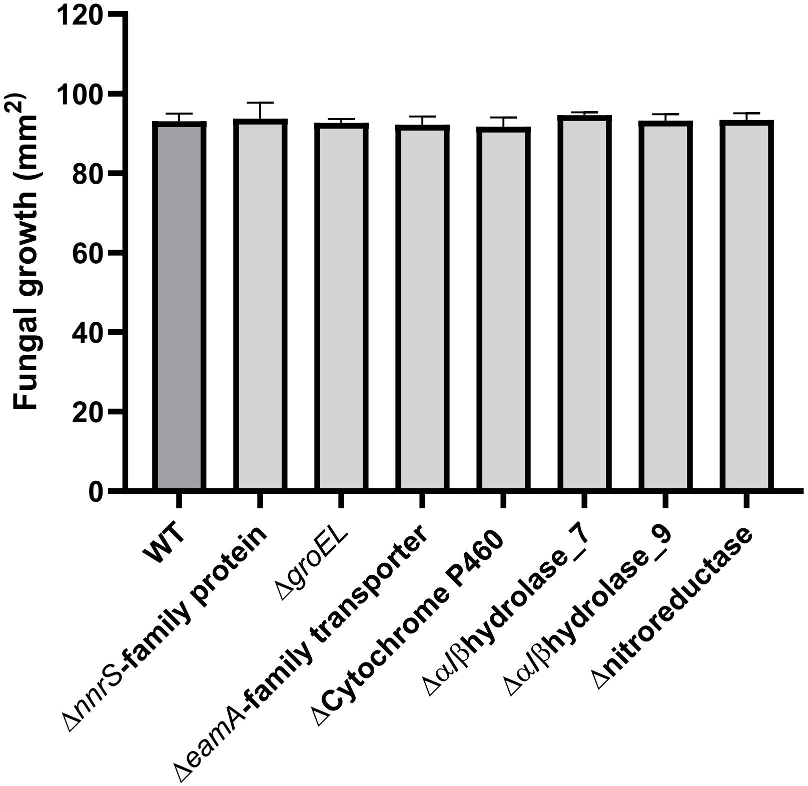


**Supplemental Figure 2:**
Genes upregulated in the Δ*hrcA P. edwinii* strain that did not have an obvious protection response phenotype. Represents three biological replicates. Error bars indicate standard deviation.

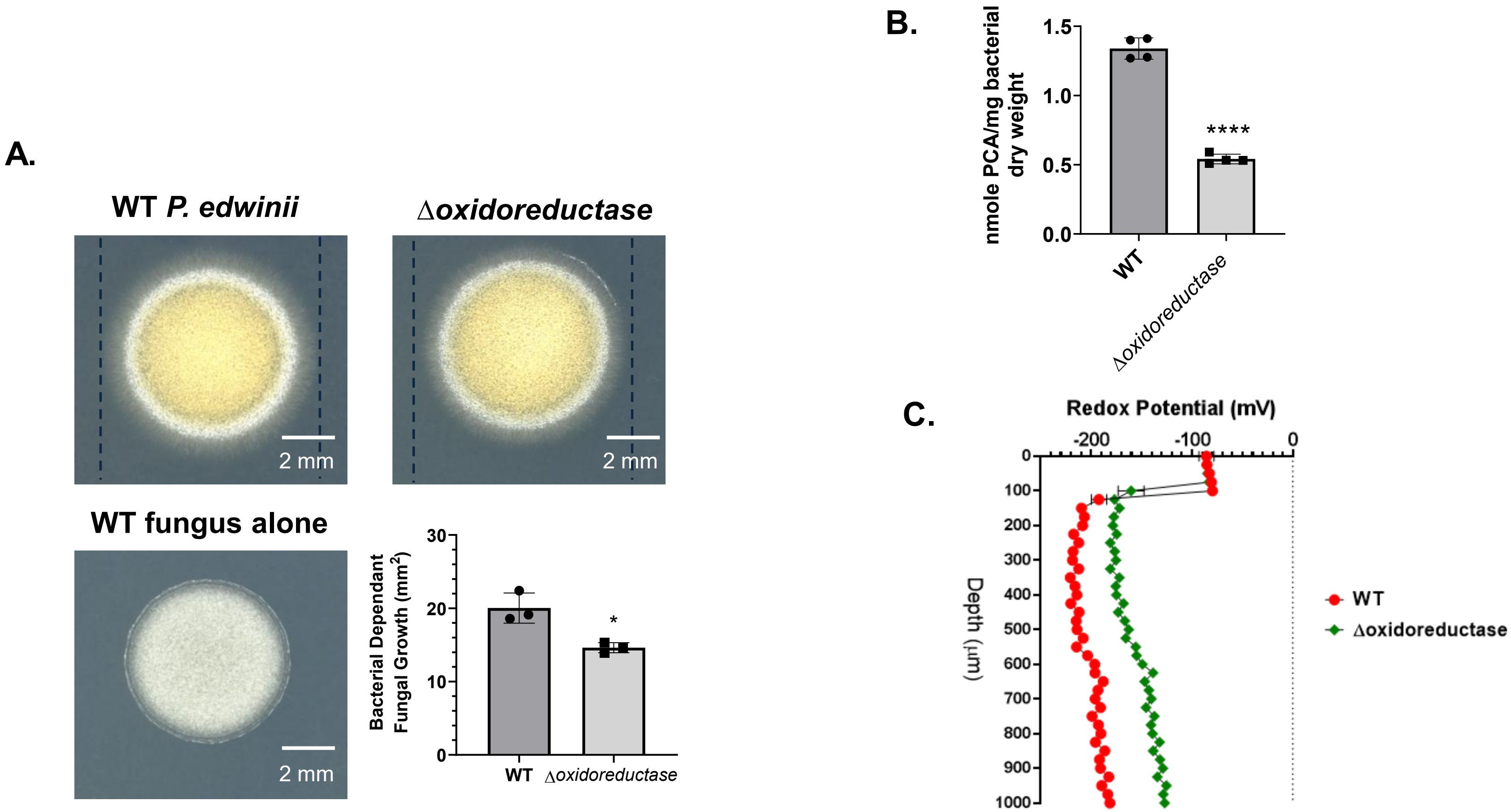


**Supplemental Figure 3:**
(A) NADP-dependent oxidoreductase (KJ38_12040) *P. edwinii* mutant has a fungal protection defect at 48hr. Dashed lines indicate diameter of wild-type co-colony. Represents three biological replicates. Scale bar indicates 2 mm, and error bars indicate standard deviation. * p-value <0.05. (B) NADP-dependent oxidoreductase mutant has a significant PCA sequestration defect at 72hr in protection activating conditions. Error bar is standard deviation. Represents four biological replicates. PCA quantified by UHPLC/MS ****p value<0.0001. (C) Redox potential (mV) through the depth of mutant bacterial colonies in protection response activating conditions at 90 hrs. Measurements were taken every 25µm with three technical measurements.

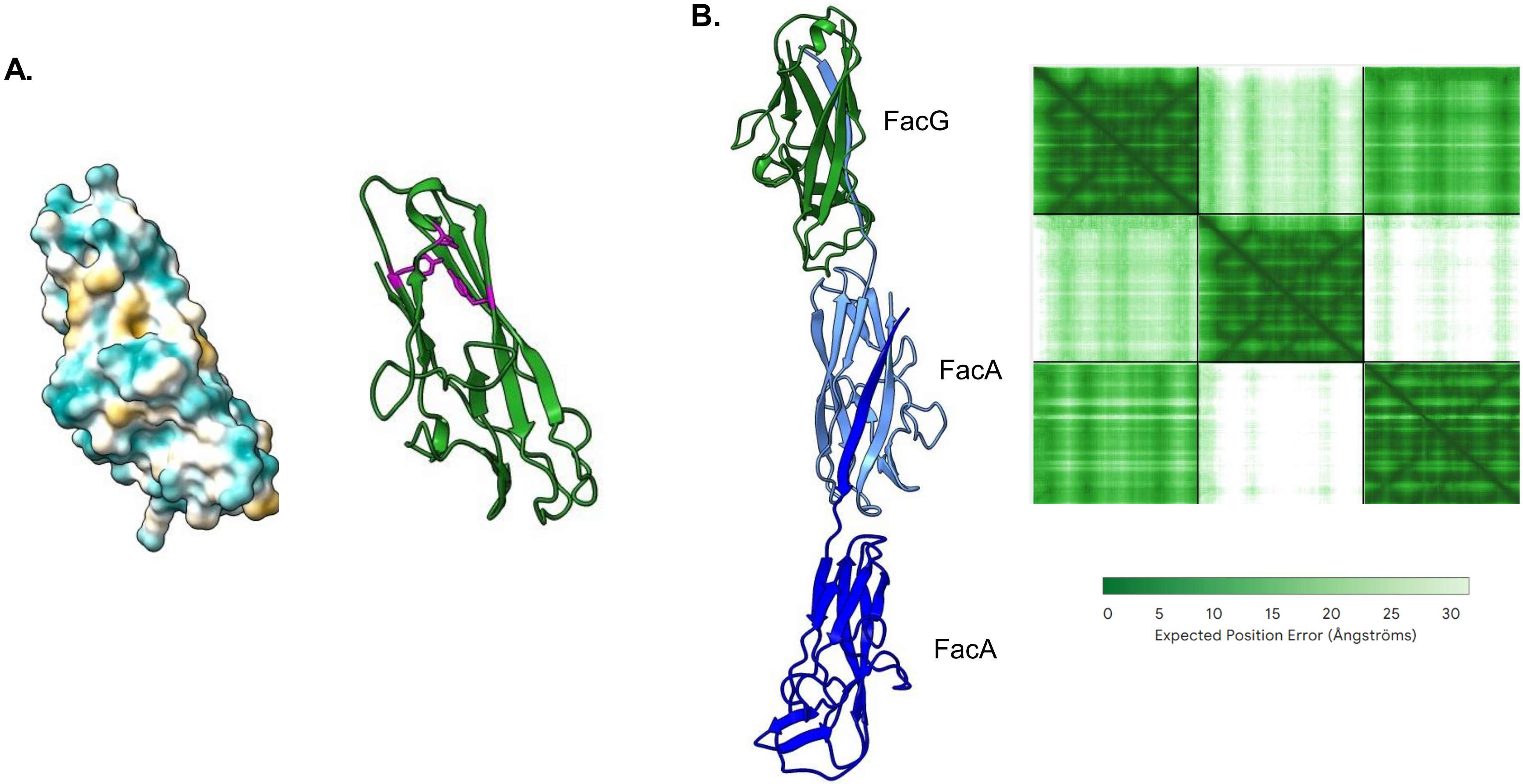


**Supplemental Figure 4:**
(A) FacG structure predictions showing hydrophobic pocket containing two Tyr (41, 104) and one Asp (93) in magenta (B) and PAE plot and ribbon structure of predicted fac polymer containing one FacG and two FacA mature proteins.

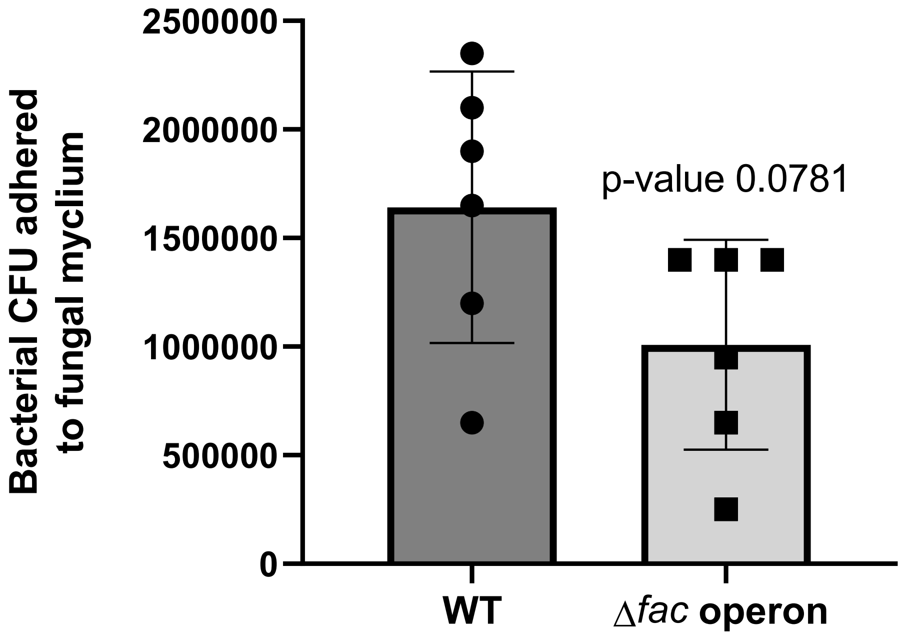


**Supplemental Figure 5:**
Bacterial CFU adhered to the fungal mycelium after 3 hours of co-incubation using the adhesion assay from Fig 3A. Represents six biological replicates. Error bars are standard deviation.

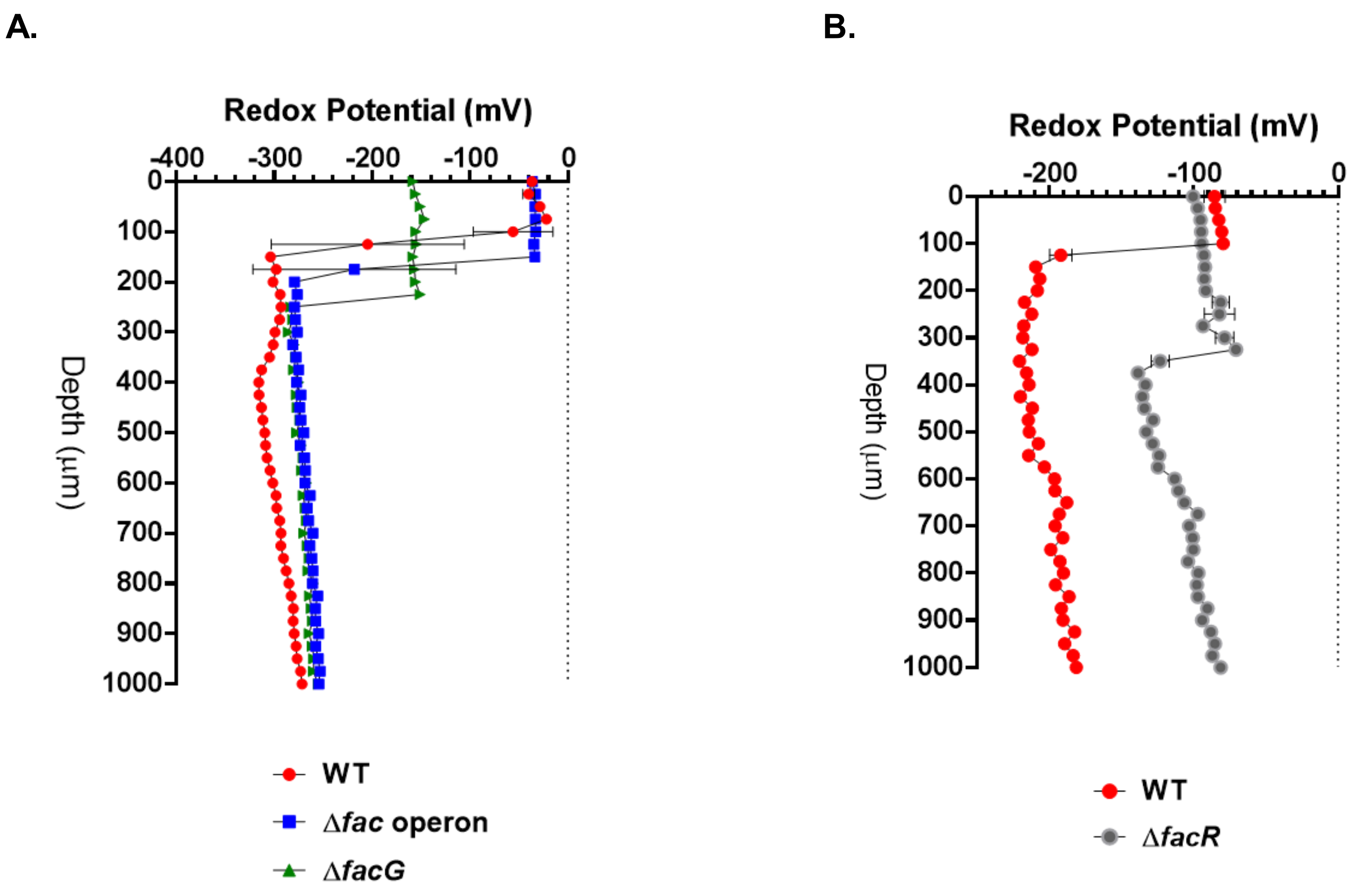


**Supplemental Figure 6:**
Biological replicates of (A) *fac* mutants' and (B) Δ*facR* redox potential measurements.

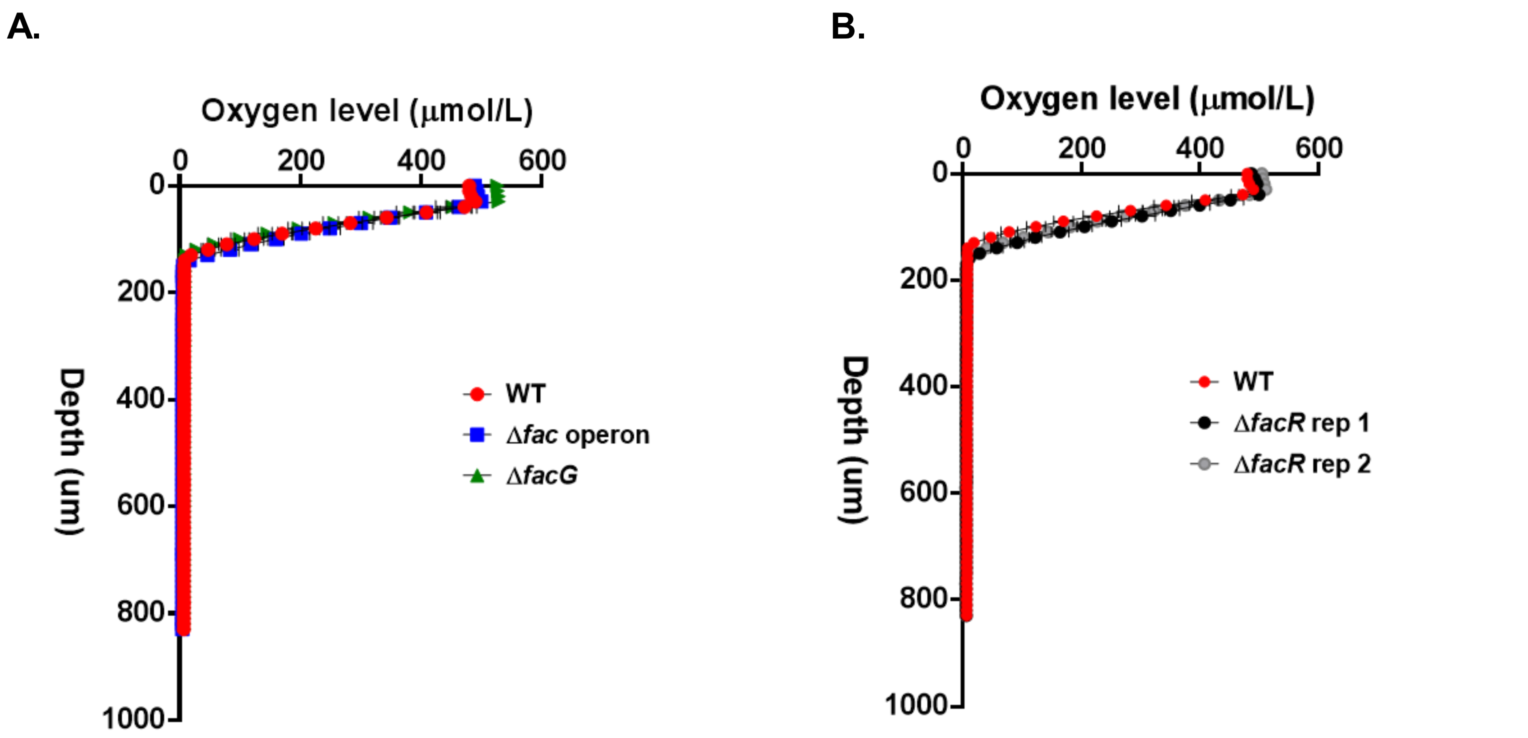


**Supplemental Figure 7:**
Biological replicates of (A) *fac* mutants' and (B) Δ*facR* oxygen level. Three technical measurements taken every 10μm.

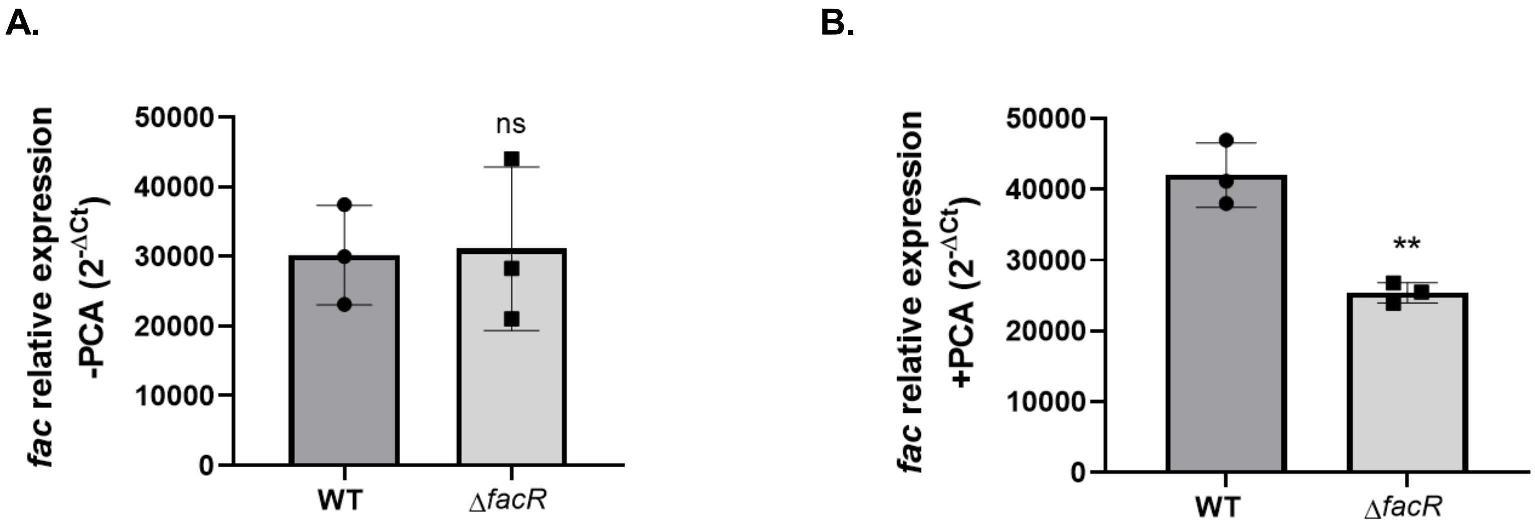


**Supplemental Figure 8:**
(A) Relative *fac* system expression in wild-type and Δ*facR*. Same data shown in Fig 8A but displayed differently. Normalized to *rpoD* expression. Units are $2^{-\Delta Ct}$ value multiplied by 0.00001 for readability. **p value<0.01.
