## Supplemental materials and methods for "A bacterial chaperone–usher pathway links interkingdom association to antimicrobial protection of a fungal partner"

| Organisms/strains | Source | Identifier | Locus Tag |
|---|---|---|---|
| *Paraburkholderia edwinii* – WT | Dahlstrom and Newman | KMD_67 | N/A |
| Paraburkholderia edwinii Δ*hrcA* | Dahlstrom and Newman | KMD_7 | KZJ38_04765 |
| Paraburkholderia edwinii Δ*facA* | This study | KMD_45 | KZJ38_12290 |
| Paraburkholderia edwinii Δ*facB* | This study | KMD_46 | KZJ38_12295 |
| Paraburkholderia edwinii Δ*facD* | This study | KMD_47 | KZJ38_12300 |
| Paraburkholderia edwinii Δ*facG* | This study | KMD_48 | KZJ38_12305 |
| Paraburkholderia edwinii Δ*facR* | This study | KMD_49 | KZJ38_15655 |
| Paraburkholderia edwinii Δ*fac* operon | This study | KMD_50 | KZJ38_12290, KZJ38_12295, KZJ38_12300, KZJ38_12305 |
| Paraburkholderia edwinii WT + pBBR1 (EV) | This study | KMD_51 | N/A |
| Paraburkholderia edwinii Δ*facG*+pBBR1(EV) | This study | KMD_52 | KZJ38_12305 |
| Paraburkholderia edwinii Δ*fac* operon +pBBR1(EV) | This study | KMD_53 | KZJ38_12290, KZJ38_12295, KZJ38_12300, KZJ38_12305 |
| Paraburkholderia edwinii Δ*facR*+ pBBR1(EV) | This study | KMD_66 | KZJ38_15655 |
| Paraburkholderia edwinii Δ*facG*+pBBR1-*facG* | This study | KMD_54 | KZJ38_12305 |

| | | | |
|---|---|---|---|
| Paraburkholderia edwinii Δ*fac* operon+pBBR1-*fac* operon | This study | KMD_55 | KZJ38_12290, KZJ38_12295, KZJ38_12300, KZJ38_12305 |
| Paraburkholderia edwinii Δ*facR*+pBBR1-*facR* | This study | KMD_56 | KZJ38_15655 |
| Paraburkholderia edwinii Δ*facR*+pBBR1-*fac* operon | This study | KMD_57 | KZJ38_15655 |
| Paraburkholderia edwinii ΔnnrS-family protein | This study | KMD_58 | KZJ38_23495 |
| Paraburkholderia edwinii Δ*groEL* | This study | KMD_59 | KZJ38_16355 |
| Paraburkholderia edwinii ΔeamA-family transporter | This study | KMD_60 | KZJ38_22480 |
| Paraburkholderia edwinii ΔCytochrome P460 | This study | KMD_61 | KZJ38_12245 |
| Paraburkholderia edwinii Δα/β hydrolase_7 | This study | KMD_62 | KZJ38_15650 |
| Paraburkholderia edwinii Δα/β hydrolase_9 | This study | KMD_63 | KZJ38_15670 |
| Paraburkholderia edwinii Δnitroreductase | This study | KMD_64 | KZJ38_12045 |
| Paraburkholderia edwinii Δoxidoreductase | This study | KMD_65 | KZJ38_12040 |
| Aspergillus calidoustus - WT | This study | KMD_21 | N/A |
| E. coli S17 λ(pir) thi pro hsdR-hsdM+ “recA RP4-2::TcMU-Km::Tn7 | Simon et al. | KMD_2 | N/A |
| **Recombinant DNA** | **Source** | **Identifier** | **Locus Tag** |
| pMQ30 | Shanks et al. | N/A | N/A |
| pMQ30 ΔfacA | This study | N/A | N/A |

| | | | |
|---|---|---|---|
| pMQ30 ΔfacB | This study | N/A | N/A |
| pMQ30 ΔfacD | This study | N/A | N/A |
| pMQ30 ΔfacG | This study | N/A | N/A |
| pMQ30 ΔfacR | This study | N/A | N/A |
| pMQ30 ΔnnrS-family protein | This study | N/A | N/A |
| pMQ30 ΔgroEL | This study | N/A | N/A |
| pMQ30 ΔeamA-family transporter | This study | N/A | N/A |
| pMQ30 ΔCytochrome P460 | This study | N/A | N/A |
| pMQ30 Δα/β hydrolase_7 | This study | N/A | N/A |
| pMQ30 Δα/β hydrolase_9 | This study | N/A | N/A |
| pMQ30 Δnitroreductase | This study | N/A | N/A |
| pMQ30 Δoxidoreductase | This study | N/A | N/A |
| pBBR1 MSC-2 | Kovach et al. | N/A | N/A |
| pBRR1 MSC-2 Rha promoter (EV) | This study | N/A | N/A |
| pBBR1 MSC-2 Rha fac operon | This study | N/A | N/A |
| pBBR1 MSC-2 Rha facG | This study | N/A | N/A |
| pBBR1 MSC-2 Rha facR | This study | N/A | N/A |

| Oligonucleotides | Source | Identifier | Locus Tag |
|---|---|---|---|
| qPCR fac operon FWD: TGATCGTCGAGGTCGTGCTG REV: TGCGATCGCAGCGACTACCA | This study | N/A | N/A |
| **Software and algorithms** | **Source** | **Identifier** | **Locus Tag** |
| Keyence VHX digital microscope measurement software | N/A | N/A | N/A |
| Google AlphaFold3 server | https://alphafoldserver.com/ | N/A | N/A |
| MEGA12 – Molecular Evolutionary Genetic Analysis | https://www.megasoftware.net/ | N/A | N/A |
| ChimeraX | https://www.cgl.ucsf.edu/chimerax/ | N/A | N/A |
| Galaxy EU biological analysis tool | https://usegalaxy.eu/ | N/A | N/A |
| R-studio | https://www.r-project.org/ | N/A | N/A |
| BioRender | https://www.biorender.com/ | N/A | N/A |
| Prism GraphPad | https://www.graphpad.com/features | N/A | N/A |
| Geneious Prime | https://www.geneious.com/ | N/A | N/A |
| SensorTrace Suite – Profiling | https://unisense.com/sensortrace-suite/ | N/A | N/A |
| SignalP | https://services.healthtech.dtu.dk/services/SignalP-6.0/ | N/A | N/A |

**Supplemental Methods:**

**Construction of bacterial strains and plasmids**

In-frame deletion strains were constructed using cloning tools adapted from *Pseudomonas* spp.(57) One kilobase upstream and downstream of the targeted gene were amplified using 5x Phusion HiFi Master Mix, with primers containing 20bp homology overhangs to a suicide plasmid (pmQ30) backbone containing a gentamicin resistance marker and *sucB* counter selection marker. pmQ30 was digested using the SmaI restriction enzyme. Plasmid backbone and PCR amplicon assembly was conducted using NEBuilder HiFi DNA Assembly master mix. Constructed cloning plasmids were transformed by electroporation into *E. coli* S17 strains and sequence confirmed. Correctly constructed plasmids were conjugated into wild-type *P. edwinii* on yeast peptone agar plates (YPA) overnight at 30C. *P. edwinii* strains containing the plasmid were selected for on PDA containing gentamicin and chloramphenicol for three days at 30C. *P. edwinii* strains that underwent homologous recombination and no longer contained the plasmid were selected for on YPA plates containing 7.5% sucrose for two days at 30C. Mutants were screened by colony PCR using Taq 2x Master Mix and confirmatory primers outside of the region of homologous recombination. Mutant strains in main figures were amplicon sequence confirmed. Expression plasmids were constructed similarly; using NEBuilder HiFi DNA Assembly master mix, a rhamnose inducible promoter system was inserted into the broad-spectrum, medium copy plasmid, pBBR1, for complementation of deleted genes. Correctly constructed complementation plasmids were electroporated into *P. edwinii* mutant or wild-type strains and transformants were selected for on Kanamycin.

**Growth conditions**

All cultures were grown in 5ml potato dextrose broth (PDB) or potato dextrose agar (PDA) plates at 30C. Liquid cultures were rotated at 70rpm and 150µM of PCA and/or L-rhamnose was added when indicated. PDA plates contained 300uM PCA and 250uM L-rhamnose for complementation or 5mM for the aggregation assay. Protection activating conditions indicate *P. edwinii* strains grown 5mm from wild-type *A. calidoustus* on PDA containing 300uM PCA. Growth time point is indicated in specific assay method section.

**RNA-sequencing**

WT and Δ*hrcA* strains of *P. edwinii* liquid cultures were normalized to 0.2 OD and 10µl were plated on potato dextrose agar (PDA) and grown at 30C for 6, 12, and 24 hours, in three biological replicates. Colonies were then scraped up with PDB, spun down at 13,600xg. RNA extraction was conducted using the Zymo Research Quick-RNA Fungal/Bacterial Miniprep kit. Extracted RNA was DNase treated using TURBO DNA-free kit, and RNA quality was confirmed through Broad Range RNA Qubit Kit and 1% gel electrophoresis stained with Sybergold. The Georgia Genomics and Bioinformatics Core performed Agilent Bioanalyzer quality check. High quality RNA was ribodepleted, library prepped, pooled, then sequenced using NextSeq 2000 (paired end, 100 cycles, 400M). RNA-seq analysis was conducted using the Galaxy EU biological analysis tool. A FastQC report was generated using the raw reads. Quality raw reads were trimmed with Trimmomatics and aligned to *P. edwinii's* reference genome using HISAT2. FeaturedCounts were conducted using *P. edwinii's* annotated reference genome, and WT and Δ*hrcA* samples were compared using DESeq2(58). Volcano plots were constructed in R-studio.

**Protein predictions and cladogram construction**

Amino acid sequences were run through SignalP to identify N-terminal signal peptide sequences that would be cleaved in mature protein. Mature amino acid sequences were run through Google AlphaFold3 server and output predicted structure CIF files were visualized in ChimeraX. CUP outer membrane usher amino acid sequences with known classification were gathered using NCBI data base and Nuccio and Baumler, 2007(24). Amino acid alignment and cladogram construction was done using MEGA12 software using maximum likelihood and adaptive bootstrap analysis.

**Protection Assay**

*P. edwinii* overnight cultures were normalized to $2.5 \times 10^{-4}$ OD in PDB. Normalized bacterial cells were mixed 1:1 with 25 million spores/mL of wild-type *A. calidoustus* spores prep. Wild-type *A. calidoustus* spores were also mixed 1:1 with 1x sterile PBS to serve as a negative control. Six µl of each co-culture or negative control were plated on PDA containing 300µM of Phenazine-1-carboxylic acid (PCA), and 250 µM L-rhamnose for co-cultures containing complement bacterial strains. At 68 or 90 hours of growth co-colonies were imaged using the Keyence VHX digital microscope. Fungal growth was quantified by measuring co-colony areas using the Keyence area analysis tool. The average area of three biological replicates of *A. calidoustus* alone were on PCA were subtracted from the co-colony area of three biological replicates to produce the area of bacterial-dependent fungal growth. Error bars indicate standard deviation and a One-way ANOVA with Dunnett's posttest was conducted for multiple comparisons or an unpaired t-test with Welch's correction for unequal variance for a single comparison to calculate statistical significance between samples (p-value).

**Bacterial CFU count**

Biological replicates of 6µl co-cultures and bacterial monocultures were grown on PDA with and without 300µM PCA for 48hr. Colonies were cut out of the agar with a sterile scalpel and put into a conical tube containing 4ml of Yeast Peptone (YP) broth. Colonies in YP were homogenized with a handheld generator probe from Fisher Scientific for 1 minute at 60% amplitude. Homogenized samples were diluted $10^{-1}$ to $10^{-5}$ in 100µl of YP. Ten µl of each co-culture dilution were spotted at the top of YP plates containing 50µg/ml of Nystatin (anti-fungal compound) and dripped down, while mono-culture samples were plated without Nystatin. Drip plates were placed in 30C for 48hr. Bacterial CFUs were counted and the total number of bacterial CFUs in the original co-culture or monoculture spot were back-calculated. Total bacterial CFUs were averaged between four biological replicates. Error bars indicate standard deviation and a One-way ANOVA with Dunnett's posttest was conducted for multiple comparisons or an unpaired t-test with Welch's correction for unequal variance for a single comparison to calculate statistical significance between samples (p-value).

**Redox potential and oxygen concentration probes**

*P. edwinii* liquid overnights were normalized to $2.5x10^{-4}$ OD in PDB. Normalized bacterial cells and 25 million spores/ml wild-type *A. calidoustus* were diluted 1:1 with 1x sterile PBS. Monocultures of the bacterial strains and wild-type *A. calidoustus* were plated 5mm apart on PDA containing 300μM PCA to induce the protection response and grown at 30C. At 90hr of growth the bacterial colonies were probed using the Unisense glass redox microelectrode paired with a reference probe (RD 403848 and REF 10-410611) and the oxygen microelectrode (OX- 10 412139). Redox measurements were taken at every 25μm for a total of 1000μm, and oxygen measurements were taken every 10μm for a total of 1000μm into the bacterial colony of two biological replicates. Three technical measurements were taken at each depth and then averaged.

**Ultra-High Performance Liquid Chromatography Mass Spectrometry (UHPLC/MS)**

*P. edwinii* strains were normalized to $2.5x10^{-4}$ OD in PDB. Normalized bacterial cells and 25 million spores/ml wild-type *A. calidoustus* were diluted 1:1 with 1x sterile PBS. Monocultures of the bacterial strains and wild-type *A. calidoustus* were plated 5mm apart on PDA containing 300μM PCA to induce the protection response and grown at 30C. At 68hr of growth the bacterial colonies were scraped up and pelleted for 1 minute. The bacterial pellets were resuspended in 200μl of sterile 1x sterile PBS and pelleted again. The bacterial pellets were placed in -80C before overnight lyophilization. The supernatant was moved into a fresh epi tube and placed in the -20C overnight to allow for larger biotic particles to sediment. After a freeze thaw cycle, 150μl of the supernatant was transferred to a 0.22μm cellulose acetate Spin X column. The column was centrifuged at 13,000xg for 1 minute and the flow-through of four biological replicates was sent for liquid chromatography mass spectrometry (UHPLC/MS) at the University of Georgias Proteomics and Mass Spectrometry facility. Samples were analyzed on a Bruker Impact II mass spectrometer interfaced with a Bruker Elute UHPLC. The UHPLC was equipped with a Waters XBridge Premier BEH C18, 3.5 μm, 21. x 50 mm column. Solvent A was 0.1% formic acid and solvent B was acetonitrile with 0.1% formic acid. After 2.0 μl of sample was injected the gradient was ramped from 5% B to 54% B in 4.6 min, 54 % B to 100% B in 1.4 min then back to 5% B in 0.5 min. The flow rate was 0.3 ml/min and the column is equilibrated for 4 minutes between runs. The effluent from the UHPLC was directed to mass spectrometer 3 minutes into the HPLC run and then diverted to waste after 6 minutes. The mass spectrometer was scanned from 100 - 650 m/z at 1 Hz using MRM to isolate the 225.06 m/z ion with a 5.0 width window and 20 eV collision energy. The capillary was set at 4500 V, nebulizer 2.5 bar and dry gas 6.0 l/min of nitrogen with a dry temperature of 200 C. Sample sets were ran three time with sample sequence scrambled between each set and the calibration standards interspersed among the samples. The data was analyzed using Bruker Compass DataAnalysis - Quant Analysis 5.1. The 207.055 m/z (±0.005 m/z) extraction ion chromatogram area was used for quantification purposes. PCA quantification was normalized to bacterial dry weight, and a two tailed test was conducted to calculate statistical significance between samples (p-value).

**Aggregation Assay**

Co-colonies prepared the same way as the protection assay, except *P. edwinii* strains are normalized to $2.5 \times 10^{-3}$ OD in PDB. Three biological replicate co-cultures were plated on PDA containing 300µM PCA and 5mM L-rhamnose for 48hr at 30C. Bacterial aggregates were imaged by the Keyence VHX digital microscope.

**Adhesion Assay**

*P. edwinii* strains and 1.25 million wild-type *A. calidoustus* spores were grown in separate 5ml of PDB overnight. *P. edwinii* strains were normalized to 0.25 OD and 50 µl of normalized cells were inoculated into the 5ml of PDB containing young wild-type *A. calidoustus* mycelium growth. *P. edwinii* and *A. calidoustus* were incubated together for 30 minutes or 3 hours at 30C, 70rpm. *A. calidoustus* mycelium was washed with 5ml of PDB twice and transferred to 500µl of fresh PDB. *A. calidoustus* mycelium was homogenized with a handheld generator probe from Fisher Scientific for 1 minute at 25% amplitude. Homogenized samples were diluted $10^0$ to $10^{-4}$ in 100µl of YP. Ten µl of each dilution were spotted at the top of YP plates containing 50µg/ml of Nystatin (anti-fungal) and dripped down. Drip plates were placed in 30C for 48hr. Bacterial CFUs were counted and the total number of bacterial CFUs in the 500µl of homogenate were back-calculated. Total number of bacterial CFUs were averaged between five or six biological replicates. Error bars indicate standard deviation and a One-way ANOVA with Dunnett's posttest was conducted for multiple comparisons or an unpaired t-test with Welch's correction for unequal variance for a single comparison to calculate statistical significance between samples (p-value).

**Backscatter Scanning Electron Microscopy (BSEM)**

For co-culture samples, *P. edwinii* strains and 1.25 million wild-type *A. calidoustus* spores were grown in separate 5ml of PDB overnight. *P. edwinii* strains were normalized to 0.25 OD and 50µl of normalized cells were inoculated into the 5ml of PDB containing young wild-type *A. calidoustus* mycelium growth. *P. edwinii* and *A. calidoustus* were incubated together for 30mins to 1.5hr at 30C, 70rpm. *A. calidoustus* mycelium was washed with 5ml of 1x sterile PBS twice and transferred to 500µl of 2% paraformaldehyde in 1x PBS. A 0.1% poly-L-lysine solution was used to affix cell suspensions to coverslips for BSEM imaging with a protocol adapted from Mazia et al.(1975)(59). The suspensions were rinsed several times in the buffer and left to incubate overnight on the coverslips. The next day, the affixed suspensions were stained with 1% $OsO_4$ for 1 hour and then chemically dehydrated with an EtOH series followed by an EtOH:HDMS series. After dehydration was complete, samples were sputter-coated with a 30 nm layer of gold-palladium and viewed on a Hitachi SU3900 SEM.

**Quantitative reverse-transcriptase PCR (qRT-PCR)**

RNA from three biological replicates was prepared in the same way as the RNA-seq samples, with samples normalized to total RNA. Reverse transcriptase (RT) and a no RT as a control reaction were conducted for each DNase treated RNA sample using SuperScript IV VILO Master Mix. Quantitative PCR was conducted on each sample's cDNA with primers designed to target the transcript of interest, with no RT reaction as control for gDNA contamination. Amplification of RpoD was used as a stable endogenous control for each sample. Target primer efficiency was confirmed by standard curve using known concentrations of cDNA. The Azure Cielo qPCR systems software calculates each reaction's CT value. Each sample's ΔCT was calculated by subtracting the experimental CT from its corresponding RpoD (endogenous controls) CT value. The ΔΔCT of each sample was calculated by subtracting the experimental sample's ΔCT from the control's ΔCT. The ΔΔCT values of all three biological replicates were converted to fold change (FC) through the formula $FC=2^{(-\Delta\Delta CT)}$. All samples FC replicates were averaged and plotted in a bar graph on prism with standard deviation. An unpaired t-test with Welch's correction for unequal variance was conducted to calculate statistical significance between samples (p-value).
